# Gene sharing during enzyme recruitment reveals distinct adaptive strategies including massive, sustained duplication without divergence

**DOI:** 10.64898/2026.02.02.703383

**Authors:** Carley Z. Reid, Diego A. R. Zorio, Brian G. Miller

## Abstract

Organisms expand their metabolism by repurposing enzymes to perform new reactions. To be repurposed, an enzyme must balance its original and new functions if both contribute to fitness. If an enzyme cannot balance its functions, another candidate may take its place. Here, we used adaptive evolution on a glucokinase-deficient *Escherichia coli* containing four promiscuous surrogates to investigate enzyme recruitment when the preferred candidate, *N*-acetyl-D-mannosamine kinase (NanK), is under selective pressure to maintain its original function. We find that NanK is still recruited to restore glycolysis under conditions requiring both functions via two distinct mechanisms that leave native activity largely unaltered. In one mechanism, small-scale gene amplification precedes the appearance of two non-synonymous mutations in *nanK* that increase glucokinase activity but have little or no effect on *N*-acetyl-D-mannosamine kinase activity. In another mechanism, recruitment occurs via amplification of a ∼1000 base pair fragment that narrowly encompasses *nanK* and reaches copy numbers as high as 127. Despite maintenance of amplification for hundreds of generations, we observe no persistent mutations in any *nanK* duplicate at the level of resolution provided by 75X whole genome sequencing coverage. Our results demonstrate that gene sharing can alter the trajectory but not necessarily prevent the recruitment of a preferred promiscuous candidate during adaptive evolution when other, seemingly equal candidates are available. Our findings also reveal that evolution by Innovation-Amplification-Divergence may only be facilitated at moderate levels of gene amplification, and hindered by massive amplification, as increased gene copy number diminishes returns of individual adaptive point mutations.

**Classification:** Evolution, Biochemistry

**Significance:** The Innovation-Amplification-Divergence model posits that single multifunctional enzymes evolve into multiple monofunctional enzymes through two events: First, selective pressure on a multifunctional enzyme leads to a duplication of the gene encoding that enzyme in an organism’s genome. Second, the duplicate gene copy can freely accumulate mutations that enhance one of the encoded enzyme’s functions while the original copy can accumulate mutations that enhance the enzyme’s other encoded function. Intriguingly, our results reveal sequence divergence only in cells that experience mild amplification, and no divergence in cells that experience massive amplification. This suggests that sequence divergence may be suppressed above a certain number of gene copies, providing a new perspective on a widely accepted theory of evolution.

## Introduction

Organisms repurpose existing enzymes to perform new functions in a process called enzyme recruitment.^1^ Comparative structural analyses suggest a pervasive role of recruitment in the formation of core metabolism, as the same enzyme scaffolds appear in diverse pathways of varying evolutionary age.^2^ In addition to its historical role in shaping central metabolism, enzyme recruitment continues to drive evolution—*Sphingobium chlorophenolicum* have recently evolved a degradation pathway for the toxic pesticide pentachlorophenol from recruited enzymes.^3^ Recruitment is often facilitated by an enzyme’s secondary, promiscuous reactions that serve as “seeds” for the evolution of a new function.^1,4^ Multicopy suppression experiments demonstrate that bacterial proteomes harbor numerous enzymes that can perform the same secondary reaction.^5–8^ Thus, candidates for enzyme recruitment are often redundant. However, simply being promiscuous does not necessarily mean an enzyme can be repurposed and integrated into a new pathway.^9^ Indeed, little is known about the specific factors that have enabled recruitment in the past, or will enable recruitment in future metabolic expansion, especially from a pool of seemingly equivalent candidates.

Our laboratory has developed a model system to systematically investigate enzyme recruitment determinants in *Escherichia coli*. The *E. coli* proteome harbors four distinct enzymes that share the same promiscuous function, but are unique in their greater organismal contexts in ways that may dictate their recruitment success.^10^ Each enzyme, *N*-acetyl-glucosamine kinase (NagK), *N*-acetyl-D-mannosamine kinase (NanK), mannofructokinase (Mak), and allokinase (AlsK) can promiscuously catalyze the first step of glycolysis, the transformation of glucose into glucose-6-phosphate.^11,12^ Previous work revealed that NanK is preferentially recruited to restore glycolysis within 100 generations of adaptive laboratory evolution (ALE).^10^ NanK’s recruitment involved nonsynonymous mutations that simultaneously enhanced the catalytic efficiency of its promiscuous reaction and damaged its primary function. This change was also reflected on the organismal level, as cells with NanK mutations demonstrated decreased fitness when grown on *N*-acetylated sugars. Thus, the recruitment of NanK required a functional trade-off between its primary and promiscuous functions.

Balancing of multiple functions is regarded as a constraint in enzyme evolution, as enhancements to one activity commonly occur at a cost to the other due to overlapping catalytic sites and residues.^13^ The extent to which one activity can evolve is limited by the changes the other activity can tolerate, which relies on both intrinsic and circumstantial factors.^13^ Several groups have performed *in vitro* and *in vivo* investigations of the evolution and mutational tolerance of multi-functional enzymes, but to our knowledge, only in organisms containing one candidate.^14–17^ Therefore, these studies do not address if the requirement to balance functions can preclude an enzyme’s recruitment when catalysis by an alternative enzyme is theoretically possible. The functional trade-off exhibited by NanK provides an intriguing opportunity to directly compare adaptation when a promiscuous enzyme is or is not required to balance both of its functions, and the supporting changes in metabolism that may aid functional balancing.

Here, we test how functional balancing alters enzyme recruitment by performing ALE on glycolytically-impaired *E. coli* in the presence of NanK’s primary and alternative substrates, *N*-acetyl-D-mannosamine and glucose. We report that NanK is recruited whether or not it must perform both of its functions, but the timeline and mechanism of recruitment are unique.^10^ Here, NanK recruitment occurs through separate mechanisms that cause minimal or no change to its primary function, including gene amplification up to 127 copies or a fixed nonsynonymous mutation that enhances NanK’s glucokinase activity while leaving the native activity unchanged. We also report that some cells develop a transient mutation that leads to upregulation of an alternative glucokinase candidate, Mak. This demonstrates that functional balancing alters, but does not necessarily prevent, recruitment of a promiscuous enzyme. Intriguingly, in instances of massive gene duplication, *nanK* copies do not harbor persistent mutations. This suggests that evolution by the widely accepted Innovation-Amplification-Divergence model may be hindered if a beneficial mutation does not appear in the early stages of amplification, before significant duplication of the parental gene suppresses the fitness benefits of mutations in any single copy.

## Results

### Adaptation under conditions requiring NanK gene sharing

NanK natively catalyzes phosphorylation of *N*-acetyl-D-mannosamine during *N*-acetyl-neuraminic (Neu5Ac) acid metabolism.^18,19^ *N*-acetyl-D-mannosamine is not produced by *E. coli* K-12 grown on glucose minimal medium.^19,20^ Therefore, under the conditions of our previous experiments, NanK was not under pressure to maintain its native function because it did not encounter its native substrate.^10^ Here, we introduced selective pressure for NanK to preserve its primary function during ALE by providing cells with glucose and *N*-acetyl-D-mannosamine. Cells were supplied with Neu5Ac, which is readily transported by the Neu5Ac: H+ symporter protein (NanT) and cleaved into *N*-acetyl-D-mannosamine and pyruvate by Neu5Ac lyase (NanA).^21^ Neu5Ac is a competent *E. coli* carbon source, and its accumulation in the cytoplasm is toxic, providing pressure for its utilization.^21^ Cells grown on 11 mM glucose and 3 mM Neu5Ac achieved the greatest increase in maximum growth rate at the lowest combined sugar concentration, so 3 mM Neu5Ac was chosen as the initial concentration for ALE (Figure S1). To identify a threshold where pressure to utilize both carbon sources was maximal, *N*-acetyl-neuraminic was reduced throughout ALE.

Prior to ALE, wild type BW25113 *E. coli* were conditioned for 500 generations on M9 minimal medium containing 11 mM glucose and 3 mM Neu5Ac.^22^ This medium-adapted strain is referred to as strain CZRLA. CZRLA cells demonstrated lag times of 3.3 ± 0.4 hr, a 2.2-fold improvement over wild-type BW25113 (7.3 ± 1.0 hr) (Figure S1). After conditioning, CZRLA cells were rendered glycolytically-deficient via deletion of *glk*, which encodes glucokinase, and *ptsH*, *ptsI*, and *crr*, which encode the phosphotransferase system (PTS) components HPr, EI, and EIIA^Glc^, respectively.^23,24^ The parental ALE strain is referred to as BW25113 Δ*glk*, Δ*pts*. The genotype of this strain was confirmed by whole genome sequencing, and its phenotype was confirmed by failure to produce colonies on glucose M9 minimal medium plates.

ALE was performed on three independent Lineages of BW25113 Δ*glk*, Δ*pts* cells. Cells were serially passaged in M9 minimal medium containing 11 mM glucose and varying concentrations of Neu5Ac for 700 generations. The concentration of Neu5Ac was reduced by 50% at generations 100 (to 1.5 mM), 300 (to 0.75 mM), and 500 (to 0.375 mM). The final 100 generations of ALE were performed at 0 mM Neu5Ac. Cells were grown to an approximate population of 4 x 10^11^ cells, then passaged to an initial population of ∼1 x 10^10^ cells to prevent loss of rare mutations.^25^ As is common in ALE studies, cells accumulated mutations throughout medium-conditioning, genetic editing, and adaptation that bear a non-obvious effect on cellular fitness (Figure S2). These are discussed in the Supplementary Information.

### Some mutational signatures of glucose processing and NanK recruitment mirror past work

Neu5Ac supplementation led to unique recruitment trajectories compared to our previous, glucose-only ALE.^10^ Within the first 100 generations of adaptation, no lineages acquired mutations related to *nan* operon regulation that were observed in past work.^10^ All Lineages instead acquired transient mutations in *cyaA* (encoding adenylate cyclase, CyaA). These were most often truncations or frameshifts, and various *cyaA* mutations appear through the rest of ALE (Figure 1). Deactivating *cyaA* mutations have been observed in other ALE experiments on *E. coli* lacking the PTS or growing on an alternative carbon source.^26,27^ Despite low levels of cAMP in PTS^-^ strains with disrupted *cyaA*, and therefore downregulation of pathways typically activated by the cAMP-crp complex, galactose-catabolism genes including *galP* were significantly upregulated in a PTS^-^strain.^26^ In addition to repression by NanR, the *nan* operon is positively regulated by crp, and thus subject to catabolite repression.^28^ However, in the absence of an intact PTS and the presence of the *nan* inducer Neu5Ac, *nan* expression is expected to be uninhibited.

**Figure 1:**
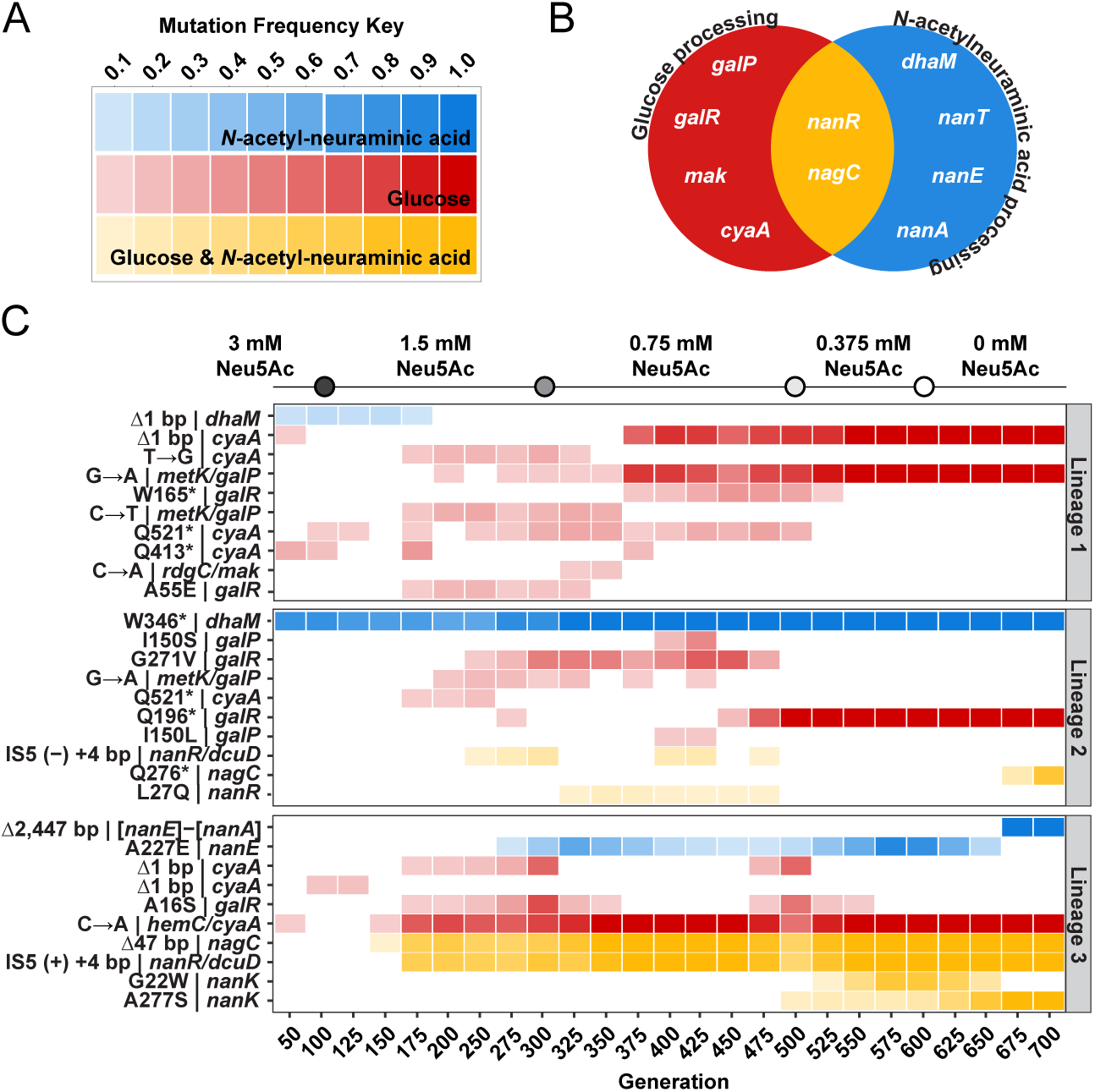
Mutations related to carbon source utilization in all replicate Lineages across adaptation. (A) Key demonstrating the relationship between mutation frequency, color opacity, and mutation effect. Only mutations that affect processing of Neu5Ac (blue), glucose (red), or both carbon sources (yellow) are shown. This classification is based on literature precedent, or on the findings of this manuscript.^19,21,26–28,32,36^ (B) A graphical depiction of mutated target genes and the classification of their effect. (C) Substrate processing mutations accrued in all Lineages between generations 50 and 700. The affected gene and resulting mutation are described. Only mutations that appear at a minimum population frequency of 0.1 and persist for at least two consecutive sequenced generations are represented. Culture conditions during ALE are presented above the heatmaps.

Mutations in galactose-processing genes, which were mutational targets in our previous study, initially appeared above 10% population frequencies starting at generations 175-250 for all Lineages, following the first reduction in Neu5Ac concentration (Figure 1).^10^ However, most of these mutations were transient, and mutations in these targets did not reach fixation until generations 500-550. Lineages 1 and 2 developed separate point mutations between *metK* and *galP*, each in a *galR* binding site roughly 67 base-pairs upstream of the *galP* transcriptional start site.^29^ *galP* encodes a galactose permease (GalP) that promiscuously transports glucose and is negatively repressed by GalR (encoded by *galR*).^29–31^ The *metK*/*galP* transition reached fixation in Lineage 1, but was lost in Lineage 2. Lineages 1 and 2 developed multiple unique transient mutations in *galR*, but only the *galR* truncation in Lineage 2 reached fixation at generation 500. Lineage 3 developed an A16S GalR mutation, which was purified from the population by generation 575. Lineage 2 also developed either an I150S or I150L mutation in the *galP* coding region, but neither reached fixation (Figure 1). Therefore, only Lineages 1 and 2 developed fixed mutations in a galactose-processing target.

Not all Lineages demonstrated unambiguous signatures of NanK recruitment from initial polymorphism analysis (Figure 1). Lineages 2 and 3 developed the same IS*5* 4-base pair insertion in the *nanR*/*dcuD* intergenic region at generations 275 and 175, respectively. *nanR* encodes the repressor (NanR) of the *nan* operon, including *nanK*.^28,32,33^ This mutation only reached fixation in Lineage 3, and was removed from the Lineage 2 population. Lineage 2 also developed a transient L27Q mutation in NanR from generations 350 to 450. Lineages 2 and 3 developed unique disruptive mutations in *nagC*. *nagC* encodes the *N*-acetyl-glucosamine repressor protein (NagC), which exhibits regulatory control on both *galP* and gene products operating downstream of NanK in Neu5Ac processing.^29,34^ A *nagC* truncation appeared at generation 675 in Lineage 2 and reached 79% frequency by generation 700. In Lineage 3, a *nagC* frameshift was present at the same frequency as the *nanR* intergenic mutation and reached fixation by generation 375. Notably, no Lineages acquired the previously-identified non-synonymous NanK mutations, L84P or L84R, across 700 generations of ALE.^10^ Additionally, Lineage 1 did not acquire any mutations in the *nan* operon or other NanK-recruitment-associated mutational targets identified in our previous study.^10^ Lineage 3 is the only population that demonstrated fixation of any mutations in the *nan* operon (Figure 1).

### Lineages acquired unique mutations in expected and unexpected loci

Intriguingly, Lineage 1 developed a fleeting C➔A transversion in the intergenic region between *rdgC* and *mak* between generations 325 and 375 (Figure 1). We previously demonstrated that this exact transversion causes an up-mutation in the *mak* promoter region that leads to a 100-fold enhancement of *mak* transcription.^12^ Lineage 3 developed several unique mutations in the *nan* operon. An A277E NanE mutation was present between generations 300 and 650, reaching a peak frequency of 88% at generation 575. NanE is an *N*-acetyl-D-mannosamine-6-phosphate epimerase and an allosteric activator of NagB, which diverts *N*-acetyl-D-glucosamine metabolism into glycolysis.^19,35^ Lineage 3 also developed fixed deletions of *nanT*, *nanE*, and *nanA* at generation 675.^19,21^ Most notably, Lineage 3 was the only population in the current study to acquire persistent, high-frequency mutations in NanK. Cells acquired either A277S or G22W NanK mutations at generations 500 or 525. A277S NanK reached fixation at generation 675 (Figure 1).

### Maximal growth rates are comparable with cells adapted without NanK functional balancing

To investigate the fitness effects of accrued mutations, growth assays on M9 minimal medium were performed for all Lineages between generations 100 and 700. Assays were performed on medium containing either 11 mM glucose or 3 mM Neu5Ac. Strain CZRLA was used as a control for wild-type glycolytic function. Previously evolved strains with L84P or L84R NanK mutations were used as a control for strains that were evolved without Neu5Ac.^10^ Average maximum growth rates on minimal medium were plotted for all strains that grew to saturation within 3 days (Figure S3). All cells reached approximately the same maximum growth rate on glucose minimal medium by generation 700 (Figure S3, Table S1). Lineage 3 cells were essentially unchanged in fitness from generations 400 to 700. In contrast, Lineages 1 and 2 experienced significant increases in maximum growth rate between generations 600 and 700. Lineage 3 maximum growth rates are statistically insignificant from cells with an L84 NanK mutation (Table S1). The Lineage 2 growth rate at generation 700 is not significantly different from cells with L84P NanK (0.22 ± 0.02 hr^-1^). Lineage 1 cells achieved their peak maximum growth rate at generation 700 but nonetheless grow more slowly than cells with L84 NanK mutations (Figure S3).

While no Lineages approached the growth rate of CZRLA cells on glucose minimal medium, Lineages 1 and 2 only fell below the CZRLA growth rate on Neu5Ac starting at generation 500 (Figure S3). Lineages 1 and 2 always exceeded cells with L84 NanK mutations in growth on Neu5Ac. Intriguingly, the maximum growth rates of Lineage 1 and 2 cells were equivalent on glucose or Neu5Ac minimal medium at generation 700. In contrast, Lineage 3 cells grew on Neu5Ac minimal medium at roughly 60% of the rate they grew on glucose at generation 700. As with growth on glucose, the growth rate of Lineage 3 cells is statistically insignificant from the growth rate of cells with L84 NanK mutations starting at generation 400 (Figure S3, Table S1).

### Glucokinase and *N*-acetyl-D-mannosamine kinase activities improved throughout adaptation

To directly assess changes in enzyme activity throughout adaptation, total glucokinase and *N*-acetyl-D-mannosamine kinase activities were determined. Specific activities were measured in whole cell lysates of population samples for each Lineage at generations 100, 300, 500, 600, and 700 (Figure 2A, Table S2). These generations correspond with changes in culture medium conditions. Specific activities were compared to lysates of CZRLA cells, which were adapted for growth on glucose and Neu5Ac medium, but have intact glycolytic function.

**Figure 2:**
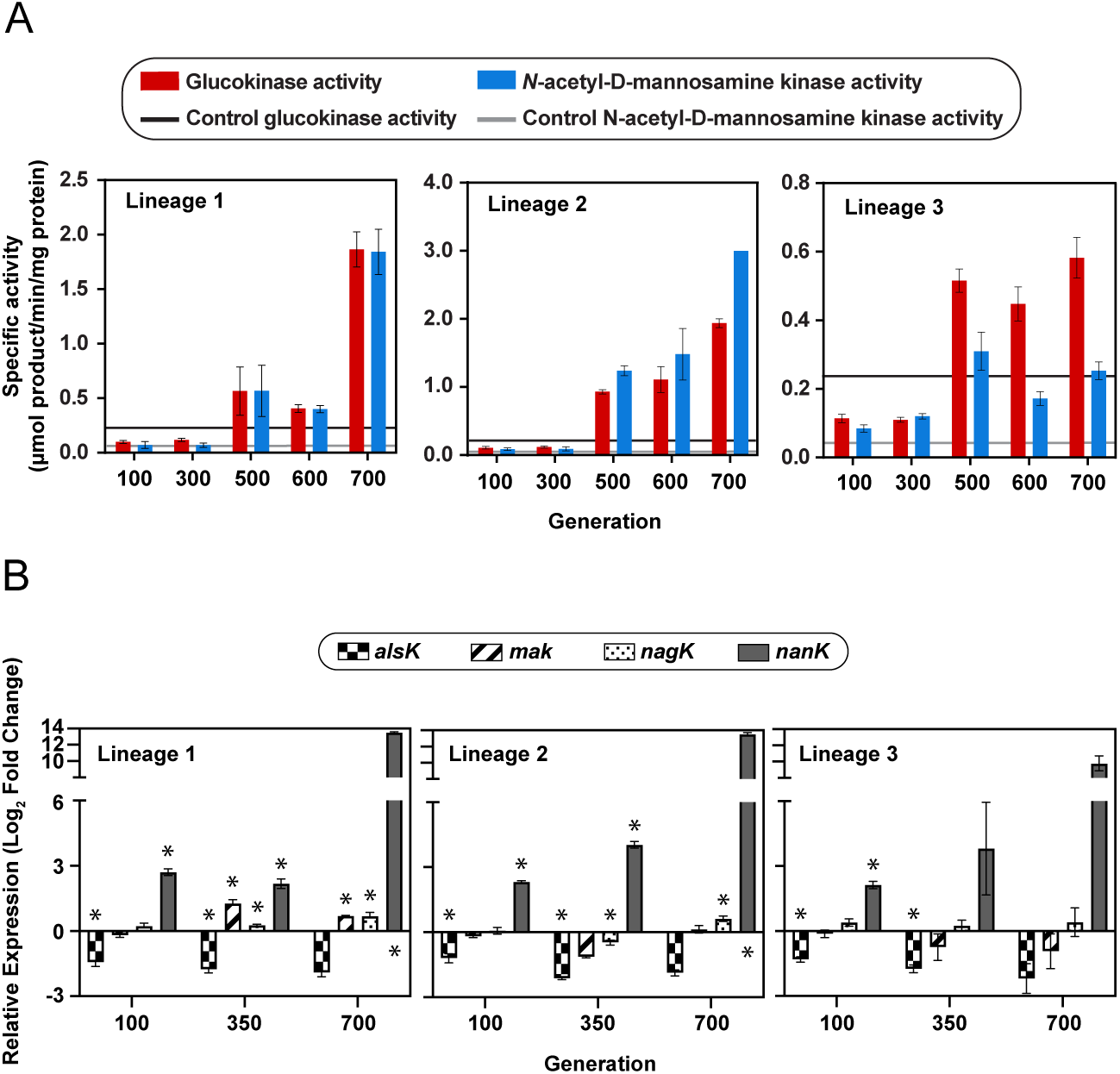
Specific glucokinase and *N*-acetyl-D-mannosamine kinase activities and transcriptional fold change in all promiscuous glucokinases throughout adaptation. (A) Specific activity measured in population cell lysates at generations 100, 300, 500, 600, and 700. Bars represent averages of 6 to 9 individual points ± SD. Glucokinase and *N*-acetyl-D-mannosamine kinase specific activities are plotted as red or blue bars, respectively. Glucokinase and *N*-acetyl-D-mannosamine kinase specific activities of CZRLA cells are represented in dark and light gray lines, respectively. (B) Relative transcription changes of promiscuous glucokinases at generations 100, 350, and 700. Data are presented as the average Log_2_ fold change in transcription, (N=3) ± SD. Relative expression was normalized to *cysG* and asterisks represent significant differences from CZRLA controls (*p < 0.05).

The average glucokinase specific activity in CZRLA lysates was 0.24 ± 0.01 µmol product min^-1^ mg protein^-1^ (Table S2). Each Lineage surpassed this value by generation 500. Lineages 1 and 3 showed comparable glucokinase activity at generation 500, at 0.57 ± 0.22 and 0.52 ± 0.03 µmol product min^-1^ mg protein^-1^, respectively, and Lineage 2 exceeded them at 0.93 ± 0.03 µmol product min^-1^ mg protein^-1^ (Figure 2A, Table S2). At generation 600, Lineages 1, 2, and 3 had specific glucokinase activities of 0.41 ± 0.04, 1.1 ± 0.1, and 0.45 ± 0.05 µmol product min^-1^ mg protein^-1^, respectively. By generation 700, Lineages 1 and 2 demonstrated equivalent glucokinase specific activity at 1.9 µmol product min^-1^ mg protein^-1^, and Lineage 3 had glucokinase activity of 0.58 ± 0.06 µmol product min^-1^ mg protein^-1^ (Figure 2A, Table S2).

*N*-acetyl-D-mannosamine kinase activity in CZRLA lysates was 0.04 ± 0.01 µmol product min^-1^ mg protein^-1^. *N*-acetyl-D-mannosamine kinase specific activity exceeded CZRLA cells for all generations and Lineages, except for Lineage 1 at generations 100 and 300 (Figure 2A, Table S4). Like glucokinase activity, the greatest increases occurred between generations 300-500 and 600-700. Lineage 1, 2, and 3 cells have *N*-acetyl-D-mannosamine kinase specific activities of 0.57 ± 0.23, 1.2 ± 0.1, and 0.31 ± 0.06 µmol product min^-1^ mg protein^-1^, respectively (Table S2). Changes in *N*-acetyl-D-mannosamine specific activity were insignificant for generations 1 and 2 between generations 500 and 600, but activity was reduced to 0.17 ± 0.02 µmol product min^-1^ mg protein^-1^ in Lineage 3. Finally, specific activity increased again for all Lineages by generation 700. Lineages 1, 2, and 3 have specific activities of 1.8 ± 0.2, 3.0 ± 0.1, and 0.25 ± 0.03 µmol product min^-1^ mg protein^-1^, respectively (Figure 2A, Table S2).

### Transcriptional analyses suggest NanK was the primary driver of adaptation

The dramatic increases in specific activity in Lineages lacking NanK mutations, as well as the transient *mak* promoter mutation in Lineage 1 (Figure 1), prompted us to investigate the extent to which additional promiscuous glucokinases participated in glucose processing during ALE. We compared the transcriptional levels of *alsK*, *mak*, *nagK*, and *nanK* in all Lineages at generations 100, 350, and 700 (Figure 2B). At generation 100, *nanK* was the only upregulated kinase out of the four candidates. Relative to CZRLA, by generation 100 *nanK* was upregulated 6.5-fold, 5-fold, and 4.4-fold for Lineages 1, 2, and 3 (Figure 2B). Conversely, *alsK* was downregulated by roughly 2.5-fold in all Lineages. The transcription of *mak* and *nagK* were unchanged from CZRLA. At generation 350, more deviations in candidate transcription occurred. Consistent with the appearance of the promoter-up mutation, *mak* was upregulated by 2.4-fold in Lineage 1 but was unchanged for other Lineages. Additionally, *nagK* was upregulated in Lineage 1 by 1.2-fold, downregulated in Lineage 2 by 1.4-fold, and unchanged in Lineage 3. *nanK* continued to be upregulated in Lineages 1 and 2 at 2.5-fold and 16.5-fold, respectively. Lineage 3 demonstrated variation in *nanK* transcription across replicates, with an average 30-fold upregulation. *alsK* remained downregulated compared to CZRLA by 3.3-to 4.4-fold. By generation 700, *alsK* was not significantly different from CZRLA for any Lineage. *mak* was upregulated by 1.6-fold in Lineage 1. *nagK* was upregulated by 1.6 and 1.5-fold in Lineages 1 and 2. *nanK* continued its trend of upregulation, with a staggering ∼12,000-fold increase in transcription in Lineages 1 and 2, and a ∼1000-fold increase in transcription in Lineage 3.

### Lineages without NanK nonsynonymous mutations developed massive amplification of the locus

To resolve the dramatic increases in both specific activity and *nanK* transcription in Lineages 1 and 2, we searched for evidence of gene duplication. Raw read coverage depth from Illumina whole genome sequencing data was used to screen for genome-wide relative copy number increases in all Lineages across 700 generations of ALE. This analysis revealed that *nanK* was dramatically amplified in Lineages 1 and 2 (Figure 3, S4). *nanK,* along with fragments of *nanE* and *nanQ*, is the only gene that is differentially amplified in comparison to strain CZRLA at generation 700 (Figure S5). Genomic copy numbers estimated from normalized Illumina read coverage depths suggest that *nanK* levels peaked at an average of 35 copies and 127 copies, respectively, for Lineages 1 and 2 at generation 700. Amplification of *nanK* was first observed at generation 350 at 11.8% population frequency in Lineage 1. By generation 375, 76.3% of the population had the amplification spanning *nanK*. The amplification persisted through the rest of adaptation, reaching a peak frequency of 97.6% at generation 700 (Figure S4).

**Figure 3:**
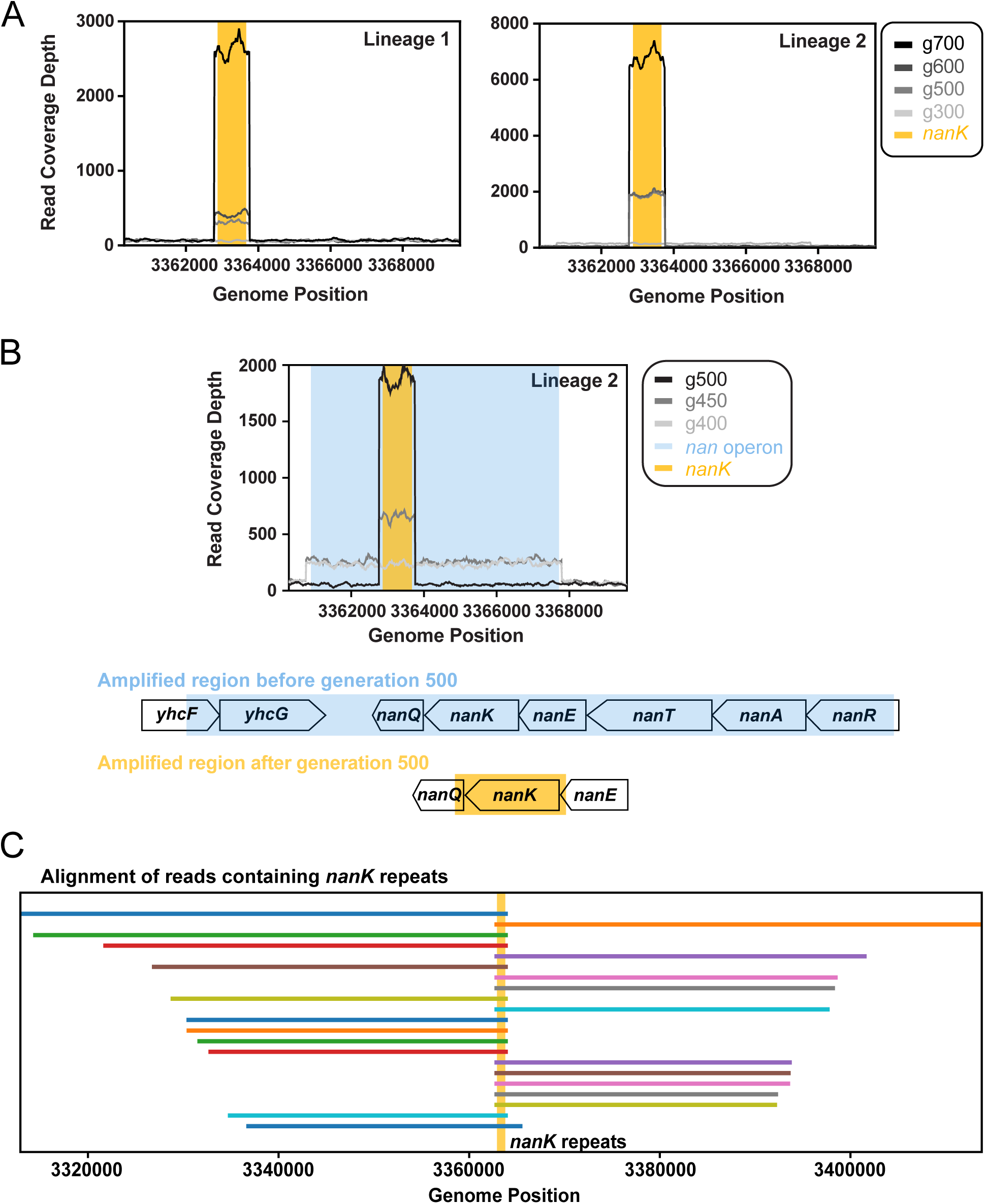
Illumina and ONT sequencing data demonstrating the amplification of *nanK* in Lineages 1 and 2. (A) Illumina raw read coverage depth of the region spanning the *nan* operon at generations 300, 500, 600, and 700 for Lineages 1 (left) and 2 (right). The region encoding *nanK* is yellow. (B) Illumina raw read coverage depth of the region spanning the *nan* operon for Lineage 2 at generations 400, 450, and 500 (top). The region encoding the *nan* operon, which was initially amplified before generation 500, is blue. The region encoding *nanK*, which is amplified after generation 500, is yellow. A graphical depiction of the genes and gene fragments included in the amplifications (bottom). (C) A representative Lineage 1 sample demonstrating alignment of ONT reads to the canonical neighboring genes of *nanK* in the *E. coli* BW25113 genome (CP009273.1). *nanK* repeats are collapsed into the *nanK* locus (highlighted vertically in yellow) for demonstrative purposes.

Unlike Lineage 1, Lineage 2 initially experienced amplification across several genes (Figures 3, S4). Amplification began at generation 250 at 51.4% population frequency and spanned the entire *nan* operon. This amplification reached a peak of 99.6% frequency at generation 450. However, an amplification spanning only *nanK* appeared in Lineage 2 in generation 425 at 7.5% frequency and reached 65.7% by generation 450. The *nan* operon amplification was ultimately lost, and the *nanK* amplification reached a peak population frequency of 99.1% at generation 700. Overlapping population frequencies of the unique amplifications suggested that a subpopulation of cells with the *nan* operon amplification initially had the *nanK* amplicon nested within it. Thus, the *nan* operon amplification narrowed to amplification of solely *nanK* from generations 475 to 700 (Figures 3, S4).

The structure of *nanK* amplification was investigated using ONT long-read sequencing on gDNA isolated from 10 clones of generation 700 populations. Five clones each were assessed from Lineages 1 and 2. Alignment of ONT reads containing *nanK* to the *E. coli* BW25113 genome revealed that *nanK* amplification occurred in its native chromosomal locus (Figure 3). This alignment also demonstrated that *nanK* repeats were complete, consecutive, and nonoverlapping. In a single ONT read, the maximum number of *nanK* repeats was 35 ± 5 for Lineage 1, and 53 ± 8 for Lineage 2. Notably, the maximum number of *nanK* repeats was limited by read length for Lineage 2 samples. Given the agreement of *nanK* copy number in Lineage 1 analyses from Illumina and ONT sequencing, it is likely that the true *nanK* copy number of Lineage 2 cells is more accurately reflected by estimates from Illumina read coverage depth (Figure S4). The amplified region spanning *nanK* includes 150 or 160 base pairs of interstitial sequence for Lineages 1 and 2, respectively. Flanking each *nanK* copy are small regions of the end of *nanE* and the beginning of *nanQ*, but no other complete genes, at generation 700 (Figure 3). Lineage 3 also demonstrated amplification, but it was smaller in magnitude and timescale. Between generations 350 and 600, *nanK* was amplified to a maximum estimated copy number of 3 based on Illumina read coverage depth. The amplification spanned *nanT, nanE,* and *nanK* and reaches a maximum frequency of 70.7% at generation 400 (Figures S4, S5).

### Unique NanK mutations enhanced its promiscuous activity with moderate impact on primary activity

Steady-state kinetic assays were performed on each NanK variant observed in Lineage 3, A277S and G22W, and wild-type NanK to investigate changes in glucokinase and *N*-acetyl-D-mannosamine kinase activity (Table 1, Figures S6, S7). Both variants have enhanced glucokinase activity compared to wild-type NanK, with improved *k*_cat_ and *k*_cat_/*K*_m_ values (Table 1). A277S NanK is the most efficient glucokinase, with a *k*_cat_/*K*_m_ of 5.0 x 10^3^ M^-1^ sec^-1^, and G22W NanK displays a *k*_cat_/*K*_m_ of 2.7 x 10^3^ M^-1^ sec^-1^ (Table 1). A277S NanK has a *k*_cat_ of 51 sec^-1^, and G22W NanK has a *k*_cat_ of 35 sec^-1^, each improved from the wild-type NanK at 23 sec^-1^. Changes in *K*_m_ are statistically insignificant for each variant (student’s t-test, p > 0.05).

**Table 1:**
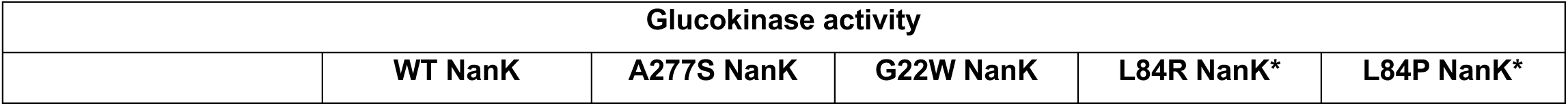
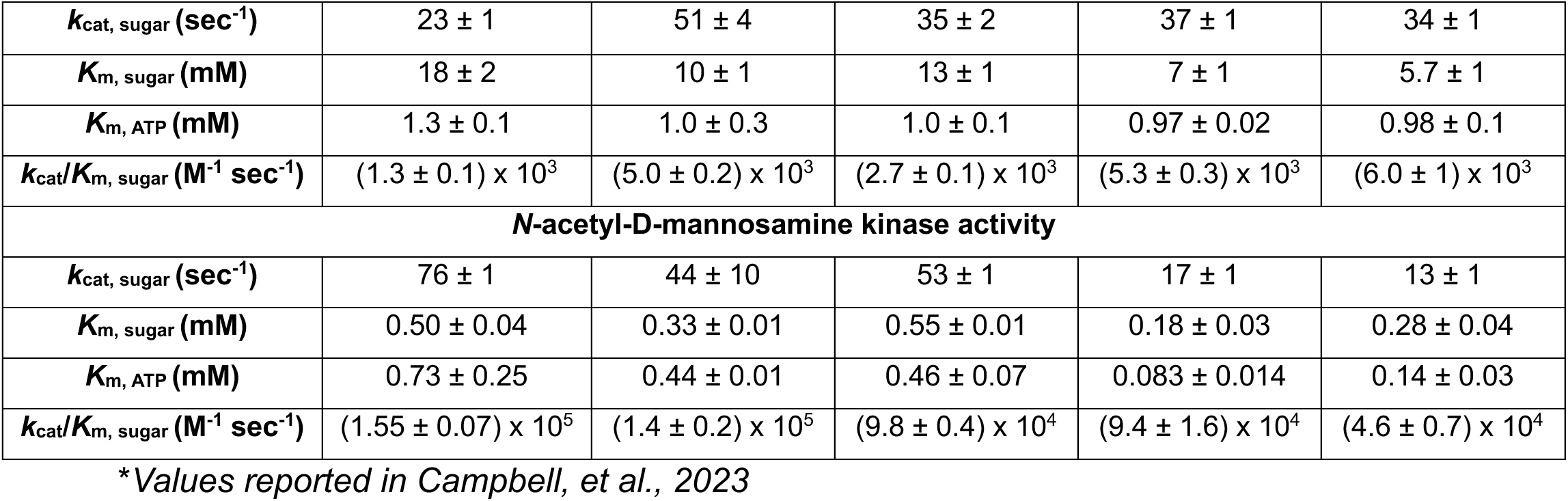
Steady-state kinetic parameters of wild type and variant NanK

| Glucokinase activity |  |  |  |  |  |
| --- | --- | --- | --- | --- | --- |
|  | WT NanK | A277S NanK | G22W NanK | L84R NanK* | L84P NanK* |
| $k_{cat, \text{ sugar }} (\text{sec}^{-1})$ | $23 \pm 1$ | $51 \pm 4$ | $35 \pm 2$ | $37 \pm 1$ | $34 \pm 1$ |
| $K_m, \text{ sugar } (\text{mM})$ | $18 \pm 2$ | $10 \pm 1$ | $13 \pm 1$ | $7 \pm 1$ | $5.7 \pm 1$ |
| $K_m, \text{ ATP } (\text{mM})$ | $1.3 \pm 0.1$ | $1.0 \pm 0.3$ | $1.0 \pm 0.1$ | $0.97 \pm 0.02$ | $0.98 \pm 0.1$ |
| $k_{cat}/K_m, \text{ sugar } (\text{M}^{-1} \text{sec}^{-1})$ | $(1.3 \pm 0.1) \times 10^3$ | $(5.0 \pm 0.2) \times 10^3$ | $(2.7 \pm 0.1) \times 10^3$ | $(5.3 \pm 0.3) \times 10^3$ | $(6.0 \pm 1) \times 10^3$ |
| ***N*-acetyl-D-mannosamine kinase activity** |  |  |  |  |  |
| $k_{cat, \text{ sugar }} (\text{sec}^{-1})$ | $76 \pm 1$ | $44 \pm 10$ | $53 \pm 1$ | $17 \pm 1$ | $13 \pm 1$ |
| $K_m, \text{ sugar } (\text{mM})$ | $0.50 \pm 0.04$ | $0.33 \pm 0.01$ | $0.55 \pm 0.01$ | $0.18 \pm 0.03$ | $0.28 \pm 0.04$ |
| $K_m, \text{ ATP } (\text{mM})$ | $0.73 \pm 0.25$ | $0.44 \pm 0.01$ | $0.46 \pm 0.07$ | $0.083 \pm 0.014$ | $0.14 \pm 0.03$ |
| $k_{cat}/K_m, \text{ sugar } (\text{M}^{-1} \text{sec}^{-1})$ | $(1.55 \pm 0.07) \times 10^5$ | $(1.4 \pm 0.2) \times 10^5$ | $(9.8 \pm 0.4) \times 10^4$ | $(9.4 \pm 1.6) \times 10^4$ | $(4.6 \pm 0.7) \times 10^4$ |

\*Values reported in Campbell, et al., 2023

G22W NanK demonstrates modest changes in its native *N*-acetyl-D-mannosamine kinase activity (Table 1). The *k*_cat_/*K*_m_ of G22W NanK is 9.8 x 10^4^ M^-1^ sec^-1^, and its *k*_cat_ is 53 sec^-1^, compared to the wild-type values of 1.55 x 10^5^ M^-1^ sec^-1^ and 76 sec^-1^. The G22W *K*_m_ is unchanged. In contrast, A277S NanK does not differ statistically from wild-type NanK in *k*_cat_, *K*_m_, and *k*_cat_/*K*_m_ (Table 1). A277S NanK has a *k*_cat_/*K*_m_ of 1.4 x 10^5^ M^-1^ sec^-1^, a *k*_cat_ of 44 sec^-1^, and a *K*_m_ of 0.33 mM.

### NanK duplicates do not demonstrate significant mutations throughout adaptation

To further investigate the extent to which *nanK* copies developed mutations in Lineages 1 and 2, all Illumina sequencing reads aligning to the *nanK* locus were investigated for base pair reads that deviated from the reference sequence at any position, generation, or frequency (Figure S8). Alternate base frequency represents the number of base reads that deviated from the reference divided by the total number of bases read across *nanK* at a given timepoint. To account for intrinsic dilution of alternate base frequency as the total number of bases read increased due to *nanK* duplication, alternate base frequencies were multiplied by the estimated *nanK* copy number per generation. In Lineage 1, the highest percentage of alternate base reads at any position in *nanK* in the population was 2.2% after copy number correction. In Lineage 2, this value was 7.9%. Neither of these mutated positions were in regions encoding the L84, A277, or G22 positions. No alternate base calls in *nanK* in any generation in Lineages 1 and 2 persisted through the rest of adaptation. Alternate base pairs persisted for 2-3 sequenced time points before exiting the population in these Lineages (Figure S8).

## Discussion

In past work, NanK was recruited to restore glycolysis in Δ*glk*, Δ*pts* cells in two primary steps: First, deactivation of the repressor NanR allowed upregulation of *nanK*.^10^ Second, nonsynonymous mutations appeared in NanK that enhanced glucokinase catalytic efficiency, but decreased *N*-acetyl-D-mannosamine kinase activity and reduced organismal fitness during growth on *N*-acetylated sugars.^10^ In the current study, NanK was still recruited to restore glycolysis. However, recruitment occurred more slowly, through unique mechanisms that caused either a point mutation or massive gene amplification of NanK, and by increasing glucokinase activity without sacrificing *N*-acetyl-D-mannosamine kinase activity.

Out of three independently evolved Lineages, only Lineage 3 acquired nonsynonymous mutations in NanK (Figure 1). These were preceded by fixed mutations related to *nan* regulation, including a frameshift in *nagC* and an insertion in the intergenic *nanR*/*dcuD* region (Figure 1). Thus, the drivers of NanK recruitment in Lineage 3 cells were superficially similar to previous work, but the mutations in NanK reported here are distinct.^10^ While the previously-observed L84R and L84P NanK variants had 6- to 7-fold greater glucokinase activity than the wild type, A277S and G22W NanK show more moderate increases in activity (Table 1).^10^ Notably, A277S NanK, which reached fixation, has *N*-acetyl-D-mannosamine kinase activity that is not statistically different from wild-type NanK. Therefore, when NanK is required to balance multiple functions during enzyme recruitment, the “winning” mutation in NanK causes a ∼4-fold increase in NanK’s promiscuous function and no change in its primary function. In contrast, the fixed mutation observed in past work, L84R, caused a 1.5-fold decrease in *N*-acetyl-D-mannosamine kinase activity and a 6.1-fold increase in glucokinase activity (Table 1).^10^ It is unclear how the A277S mutation confers a significant change in glucokinase activity with no change to NanK’s native function. Both A277 and G22 are located > 10 Å from the NanK active site (Figure S9). Nonetheless, our study indicates that mutations occurring outside of the active site may provide an adaptive pathway to a promiscuous enzyme that circumvents functional trade-offs.

Unlike Lineage 3, Lineage 1 and 2 cells developed neither persistent, high-frequency nonsynonymous mutations in NanK, nor fixed mutations related to *nan* operon regulation. In fact, Lineage 1 cells never acquired mutations in any member of the *nan* operon throughout adaptation (Figure 1). Instead, these Lineages adapted through massive, sustained gene amplification up to 127 copies of a ∼1000 base-pair region including *nanK* (Figure 3). While gene amplification is a common adaptive strategy in *E. coli*, amplification is typically transient.^37,38^ In the absence of obvious polymorphisms, the amplifications in Lineages 1 and 2 appear to explain trends in glucokinase and *N*-acetyl-D-mannosamine kinase specific activities, and *nanK* relative transcription across adaptation (Figure 2). The onset of amplification coincided with initial leaps in specific activity between generations 300 and 500. Amplification and specific activity are stagnant from generations 500 to 600, and the greatest increases in both amplification and specific activities occurred between generations 600 and 700 (Figures 2A, Figure 3). Furthermore, Lineage 2 cells demonstrated earlier increases in each specific activity, and greater total *N*-acetyl-D-mannosamine kinase activity, consistent with greater average *nanK* amplification in these cells. The transitions between generations also coincided with changes in Neu5Ac concentration that are expected to increase selective pressure. At generation 500, Neu5Ac concentration is reduced to 0.75 mM, which is near the estimated *K*_m_ of Neu5Ac transport in *E. coli*, and therefore is likely the first generation where Neu5Ac transport becomes limited.^21^

By generation 700, Lineages 1 and 2 each demonstrate ∼12,000-fold transcriptional upregulation of *nanK* compared to CZRLA controls, despite complete removal of the *nan* operon inducer. Lineage 3 experiences ∼1000-fold upregulation of *nanK* at generation 700 (Figure 2B). In past work, Δ*glk,* Δ*pts* cells evolved without Neu5Ac showed initial upregulation of *nanK* by ∼4- to 8-fold at generation 10.^10^ By generation 100, following onset of NanK mutations, Lineages showed either a downregulation or reduction in upregulation in *nanK*.^10^ While it is unclear if *nanK* was ever upregulated beyond 8-fold between generations 10-100, the level of *nanK* upregulation in the current study far exceeded what was observed in past work, even in cells with the A277S NanK mutation. Cells in past work also exhibited upregulation of the alternative promiscuous candidate *alsK*, which was consistently downregulated here.^10^ Another unique attribute of this work is the appearance of a transient mutation in the *mak* promoter in Lineage 1 (Figure 1). Despite only reaching 13% frequency in the population before its removal, this promoter-up mutation has been demonstrated to cause a 100-fold increase in *mak* transcription.^12^ Indeed, Lineage 1 is the only Lineage to demonstrate upregulation of *mak* throughout adaptation. The *mak* promoter mutation represents the only mutation observed in our cumulative ALE efforts that is unambiguously related to increasing flux through a glucokinase candidate other than NanK.^10^ This implies that gene sharing broadened the scope of sampled candidates, but that ultimately, NanK is still the preferred recruitment candidate in our system. Further investigation into the specific attributes of NanK that facilitate its recruitment, or the attributes that constrain other candidate glucokinases from recruitment, may reveal why NanK is recruited despite its functional constraint.

Despite comparably low specific activities and *nanK* transcription, Lineage 3 cells demonstrated the greatest organismal fitness when grown on glucose minimal medium as early as generation 400 (Figure S3, Table S1). By generation 500, Lineage 3 maximum growth rates are statistically insignificant from cells with an L84 NanK mutation (Figure S3). Thus, functional balancing led to a mutation in a recruited enzyme that caused less enhancement of its promiscuous function, no change to its primary function, and yet the same fitness as cells evolved without pressure for functional balancing.^10^ Compared to cells evolved without Neu5Ac, Lineage 3 demonstrated additional high-frequency mutations in *nagC* and *nanE* that may contribute to this observation. Lineages 1 and 2 only approached the fitness of cells with L84 NanK mutations when grown on glucose minimal medium at generation 700. This implies that massive duplication of NanK was required to achieve the approximate fitness of strains that simply have a nonsynonymous mutation in NanK. Despite minimal or no damage to NanK, the maximum growth rates of all cells were diminished on Neu5Ac minimal medium by generation 700 (Figure S3). Lineage 3 cells have the lowest growth rate on Neu5Ac starting at generation 400, preceding any NanK mutations. Intriguingly, Lineage 3 cells at generation 700 were still able to grow on Neu5Ac despite a deletion spanning *nanT*.^32,39,40^

In the absence of gene sharing, *nanK* amplification was not observed as a prerequisite for *nanK* recruitment in three separate genetic backgrounds and ALE experiments.^10^ Instead, mutations in *nanK* appeared after a minimum of 50 generations of ALE, with no evidence of gene amplification preceding those mutations.^10^ The moderate amplification observed in Lineage 3 cells preceding two nonsynonymous mutations in *nanK* represents the only example of consecutive amplification and divergence in our experimental system (Figure S4). Cumulatively, we have identified 4 separate nonsynonymous mutations in NanK that are evolutionarily accessible and can increase glucokinase activity up to 7-fold. Yet, we fail to see a sweep of any one of these variants, nor any other mutations in *nanK*, in any of the 35-127 copies of *nanK* encoded in Lineages 1 and 2. Previous reports suggest that gene amplification does not necessarily facilitate the accumulation of beneficial point mutations, and can even hinder sequence divergence entirely under some conditions.^41,42^ Tomanek and Guet investigated the adaptive strategies employed in *E. coli* grown on galactose to increase the transcription of *galK.* They found that when galactose concentrations were low, or its initial promoter sequence was mutated to cause intrinsically higher *galK* expression, amplification of *galK* or point mutation in its promoter region were mutually exclusive. When galactose concentrations were intermediate or high, amplification and point mutation were often observed in combination.^42^ An intriguing deviation of our study from the work of Tomanek and Guet is the observation of differing adaptive strategies—namely, massive amplification without divergence or moderate amplification with divergence under the exact same conditions. Despite the prevailing wisdom of the Innovation-Amplification-Divergence model, which posits that promiscuous enzymes evolve through consecutive amplification and sequence divergence, *nanK* amplification and mutation appear to be decoupled in our experimental system.^4,10^ We do not see evidence of any “competing” subpopulations containing *nanK* mutations in Lineages 1 and 2, and amplifications quickly rise to a majority frequency once they appear in those Lineages. Thus, clonal interference does not appear to explain the lack of *nanK* mutations in those Lineages.

We posit that evolution by the Innovation-Amplification-Divergence model may be hindered by sustained and increasing amplification prior to the initial emergence of beneficial point mutations. That is, the fitness increase resulting from a beneficial *nanK* mutation will be higher if 1/3 copies bears the beneficial mutation. However, as the number of *nanK* copies increases, the weighted effect of enhanced catalytic efficiency in any one copy is diminished as this ratio decreases to 1/35 or 1/127 wild-type copies, as is observed in our system (Figure 4). Notably, a similar scenario has been observed during recruitment of γ-glutamyl phosphate reductase to serve a new function in arginine biosynthesis.^16^ From 8 total ALE Lineages, all of which showed evidence of gene amplification, only the lineage with the lowest apparent copy number accumulated a beneficial mutation that lead to sequence divergence and de-amplification. Therefore, at a moderate level of amplification, a beneficial mutation can contribute to fitness and sweep the population, but when copy numbers exceed a certain threshold, even beneficial mutations may be “muted” in their effect and lost to drift.

**Figure 4:**
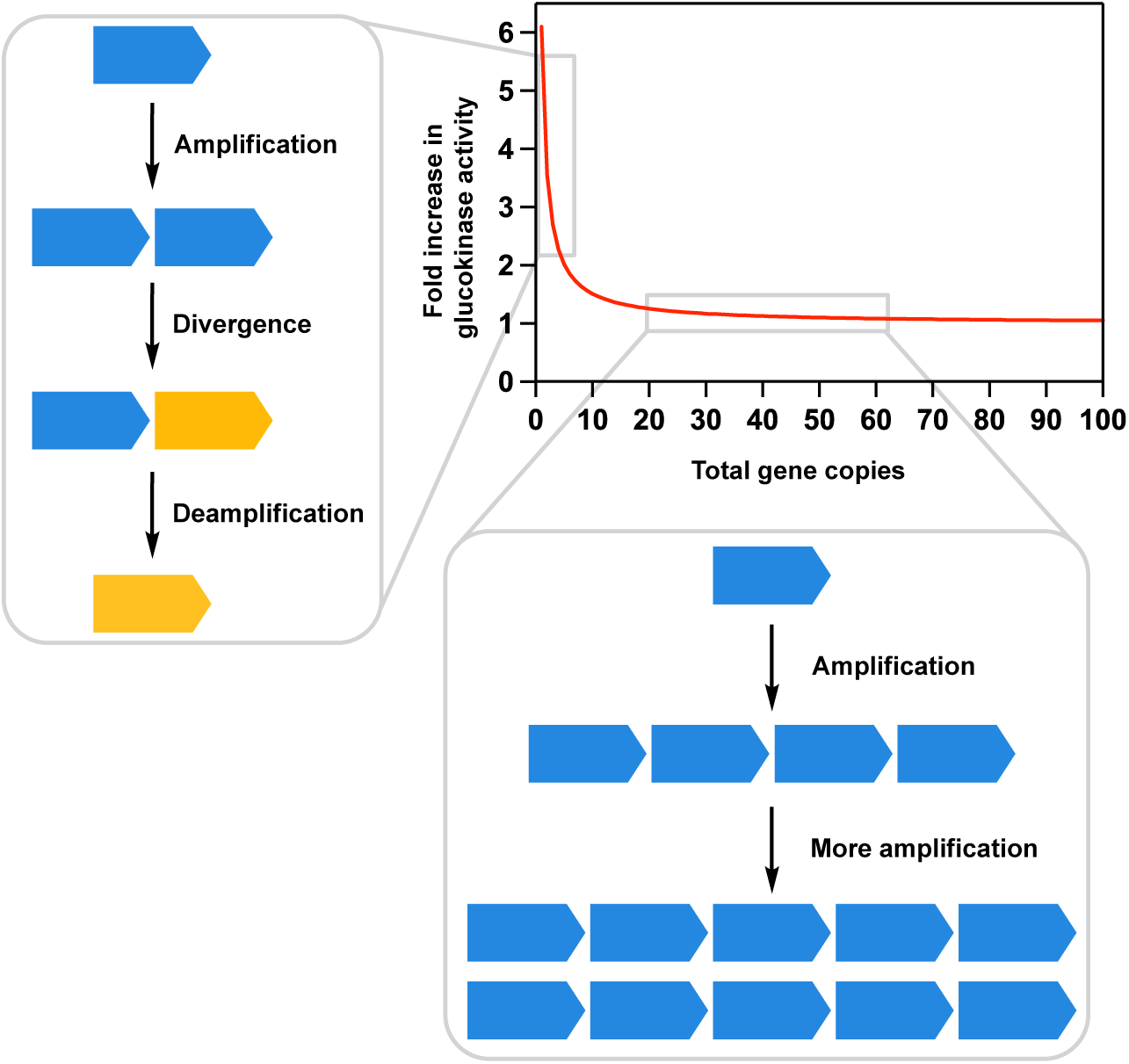
Illustration of the decreased fitness benefit of a point mutation as the copy number of an amplified gene increases. To model this effect, we used kinetic data of wild-type NanK compared to the best identified variant, L84R.^10^ At low copy number, the fold increase in total glucokinase activity when one copy has a NanK mutant is high, and this facilitates evolution that follows the Innovation-Amplification-Divergence model (left). When total copy number of *nanK* is high, fold change in total activity from one mutant NanK copy approaches 1, and *nanK* is further amplified without divergence (bottom).

## Materials and Methods

Materials and methods are described in the accompanying Supplementary Information.

## Acknowledgements

Research reported in this publication was supported by the National Institute of General Medical Sciences of the National Institutes of Health under Award Numbers R01GM133843 and R01GM157172 (B.G.M.) The content is solely the responsibility of the authors and does not necessarily represent the official views of the National Institutes of Health.

