## Supplementary Information for "Gene sharing during enzyme recruitment reveals distinct adaptive strategies including massive, sustained duplication without divergence"

**Table of Contents:**

**Supplementary Figures and Tables Referenced in Main Text** ***pg.* 3-17**

Figure S1 *pg.* 3

Figure S2 *pg.* 4-5

Figure S3 *pg.* 6

Table S1 *pg.* 7

Table S2 *pg.* 8

Figure S4 *pg.* 9-10

Figure S5 *pg.* 11-12

Figure S6 *pg.* 13

Figure S7 *pg.* 14

Figure S8 *pg.* 15-16

Figure S9 *pg.* 17

**Supplementary Results and Discussion: Additional Mutations** ***pg.* 18-22**

Figure S10 *pg.* 21

Figure S11 *pg.* 22

**Materials and Methods** ***pg.* 23-33**

Table S3 *pg.* 32

Table S4 *pg.* 33

**References** ***pg.* 34-37**

### Supplementary Figures and Tables referenced in Main Text

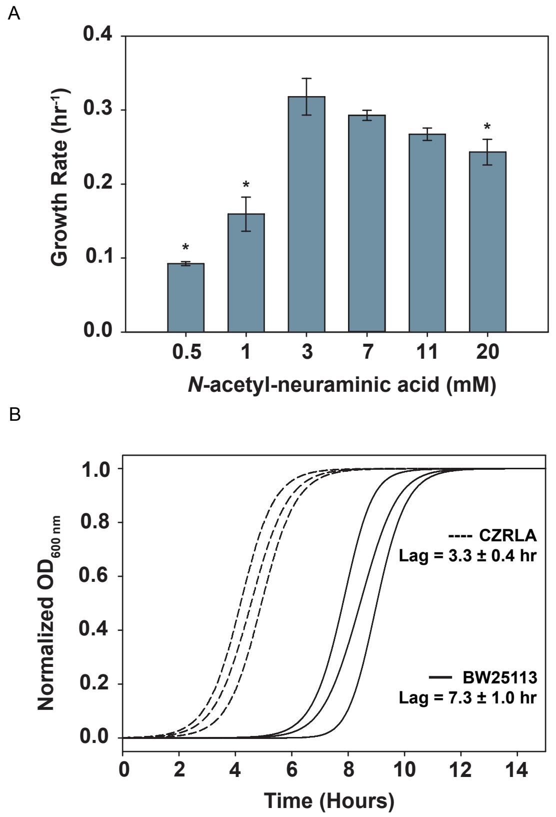


**Figure S1:** Preliminary growth tests of parental strains. (A) Maximum growth rate of BW25113 *E. coli* (previously adapted to growth on glucose minimal medium[1]) on varying concentrations of Neu5Ac. Asterisks designate concentrations where maximum growth rates are significantly different from 3 mM, as determined by a Student's t-test ($p < 0.05$) Averages are (N=3) ± SD. (B) Normalized growth curves for wild type BW25113 cells (solid line) and CZRLA cells (dashed line) grown on minimal medium containing 3 mM Neu5Ac and 11 mM glucose. Averages are (N=3) ± SD.

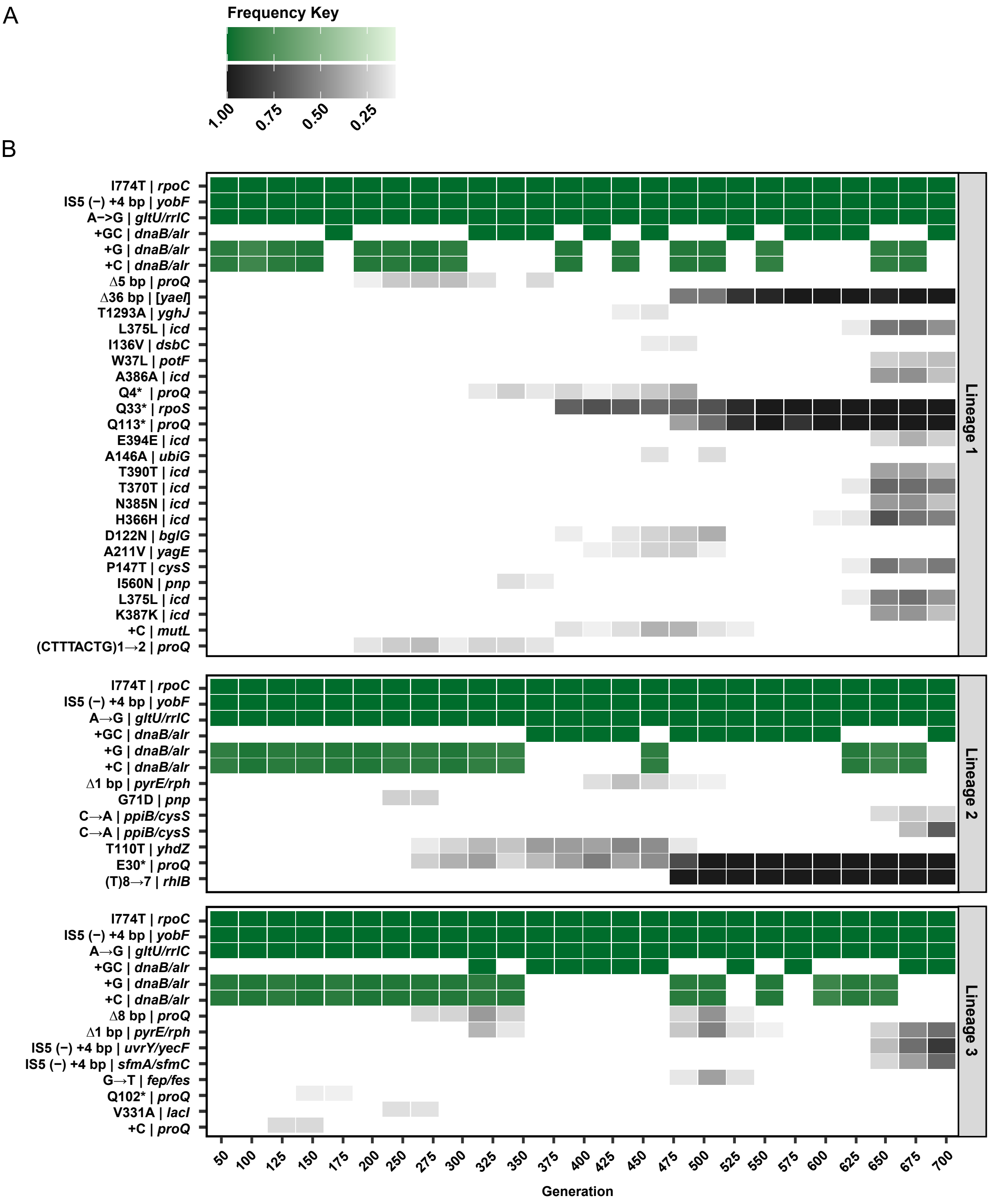


**Figure S2:** Heatmap of mutations accumulated in all Lineages that are not directly related to processing of glucose and/or Neu5Ac. (A) Mutation Frequency Key relating mutation frequency and color opacity. (B) Mutations accumulated in all Lineages that reached a minimum frequency of 0.1 and persisted for at least two consecutive sequencing timepoints. Mutations that are shared between all Lineages are presented in green. Otherwise, mutations are presented in grayscale.

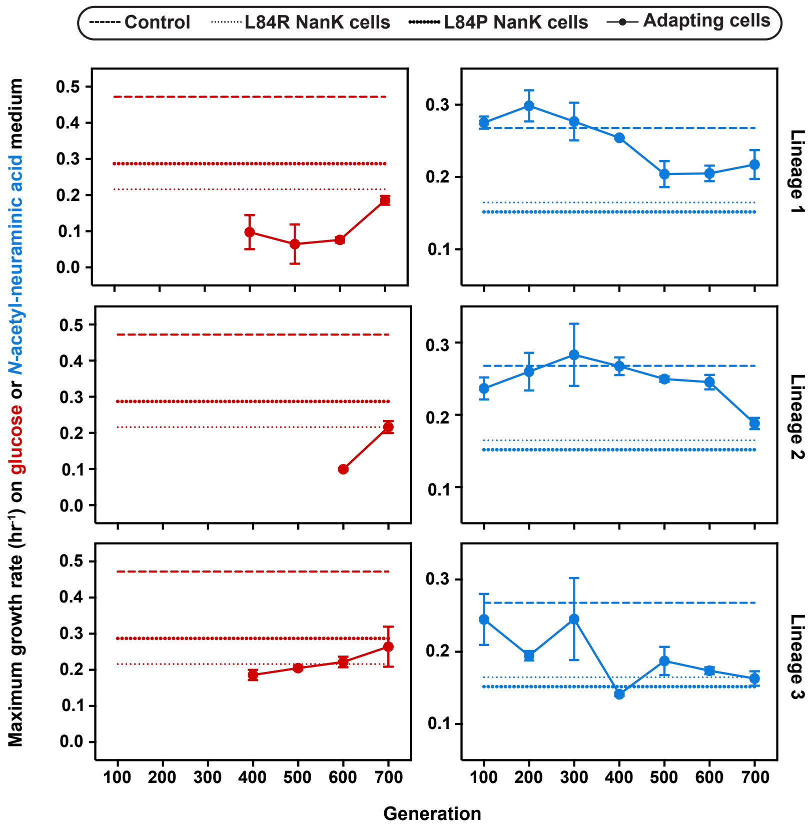


**Figure S3:** Fitness assays of replicate Lineages on minimal medium containing either glucose (11 mM) or Neu5Ac (3 mM). Maximum growth rates for cells grown on glucose minimal medium are shown in red (left). Maximum growth rates for cells grown on Neu5Ac minimal medium are shown in blue (right). Control cells (dashed line) are strain CZRLA. Adapting cells (solid line) between generations 100 and 700 are compared to BW25113 Δ<u>*glk*</u> Δ*pts* cells with either an L84R (small dotted line) or L84P (large dotted line) NanK mutation.[1]

**Table S1: Average maximum growth rates on glucose or Neu5Ac medium**

| Strain | Average max growth rate ($hr^{-1}$) on glucose minimal medium | Average max growth rate ($hr^{-1}$) on Neu5Ac minimal medium |
|---|---|---|
| CZRLA | 0.47 ± 0.03 | 0.27 ± 0.01 |
| L84P NanK | 0.29 ± 0.01 | 0.15 ± 0.03 |
| L84R NanK | 0.22 ± 0.01 | 0.16 ± 0.02 |
| CR1-100 | N/A | 0.24 ± 0.02 |
| CR1-200 | N/A | 0.26 ± 0.03 |
| CR1-300 | N/A | 0.28 ± 0.04 |
| CR1-400 | 0.10 ± 0.05 | 0.27 ± 0.01 |
| CR1-500 | 0.07 ± 0.06 | 0.25 ± 0.01 |
| CR1-600 | 0.08 ± 0.01 | 0.25 ± 0.01 |
| CR1-700 | 0.19 ± 0.01 | 0.19 ± 0.01 |
| CR2-100 | N/A | 0.28 ± 0.01 |
| CR2-200 | N/A | 0.30 ± 0.02 |
| CR2-300 | N/A | 0.28 ± 0.03 |
| CR2-400 | N/A | 0.25 ± 0.01 |
| CR2-500 | N/A | 0.20 ± 0.01 |
| CR2-600 | 0.10 ± 0.01 | 0.21 ± 0.01 |
| CR2-700 | 0.22 ± 0.02 | 0.22 ± 0.02 |
| CR3-100 | N/A | 0.28 ± 0.04 |
| CR3-200 | N/A | 0.19 ± 0.01 |
| CR3-300 | N/A | 0.25 ± 0.06 |
| CR3-400 | 0.18 ± 0.01 | 0.14 ± 0.01 |
| CR3-500 | 0.20 ± 0.01 | 0.19 ± 0.02 |
| CR3-600 | 0.22 ± 0.02 | 0.17 ± 0.01 |
| CR3-700 | 0.26 ± 0.06 | 0.16 ± 0.01 |

**Table S2: Specific activities in cell lysates throughout adaptation**

| Lineage | Generation | Specific glucokinase activity (μmol product $min^{-1}$ mg $protein^{-1}$) | Specific *N*-acetyl-D-mannosamine kinase activity (μmol product $min^{-1}$ mg $protein^{-1}$) |
|---|---|---|---|
| CZRLA | N/A | 0.24 ± 0.01 | 0.04 ± 0.01 |
| 1 | 100 | 0.10 ± 0.01 | 0.07 ± 0.03 |
| | 300 | 0.12 ± 0.01 | 0.07 ± 0.02 |
| | 500 | 0.57 ± 0.22 | 0.57 ± 0.23 |
| | 600 | 0.41 ± 0.04 | 0.40 ± 0.03 |
| | 700 | 1.9 ± 0.2 | 1.8 ± 0.2 |
| 2 | 100 | 0.11 ± 0.02 | 0.09 ± 0.2 |
| | 300 | 0.12 ± 0.01 | 0.09 ± 0.3 |
| | 500 | 0.93 ± 0.03 | 1.2 ± 0.1 |
| | 600 | 1.1 ± 0.1 | 1.5 ± 0.4 |
| | 700 | 1.9 ± 0.1 | 3.0 ± 0.1 |
| 3 | 100 | 0.11 ± 0.01 | 0.08 ± 0.01 |
| | 300 | 0.11 ± 0.01 | 0.12 ± 0.01 |
| | 500 | 0.52 ± 0.03 | 0.31 ± 0.06 |
| | 600 | 0.45 ± 0.05 | 0.17 ± 0.02 |
| | 700 | 0.58 ± 0.06 | 0.25 ± 0.03 |

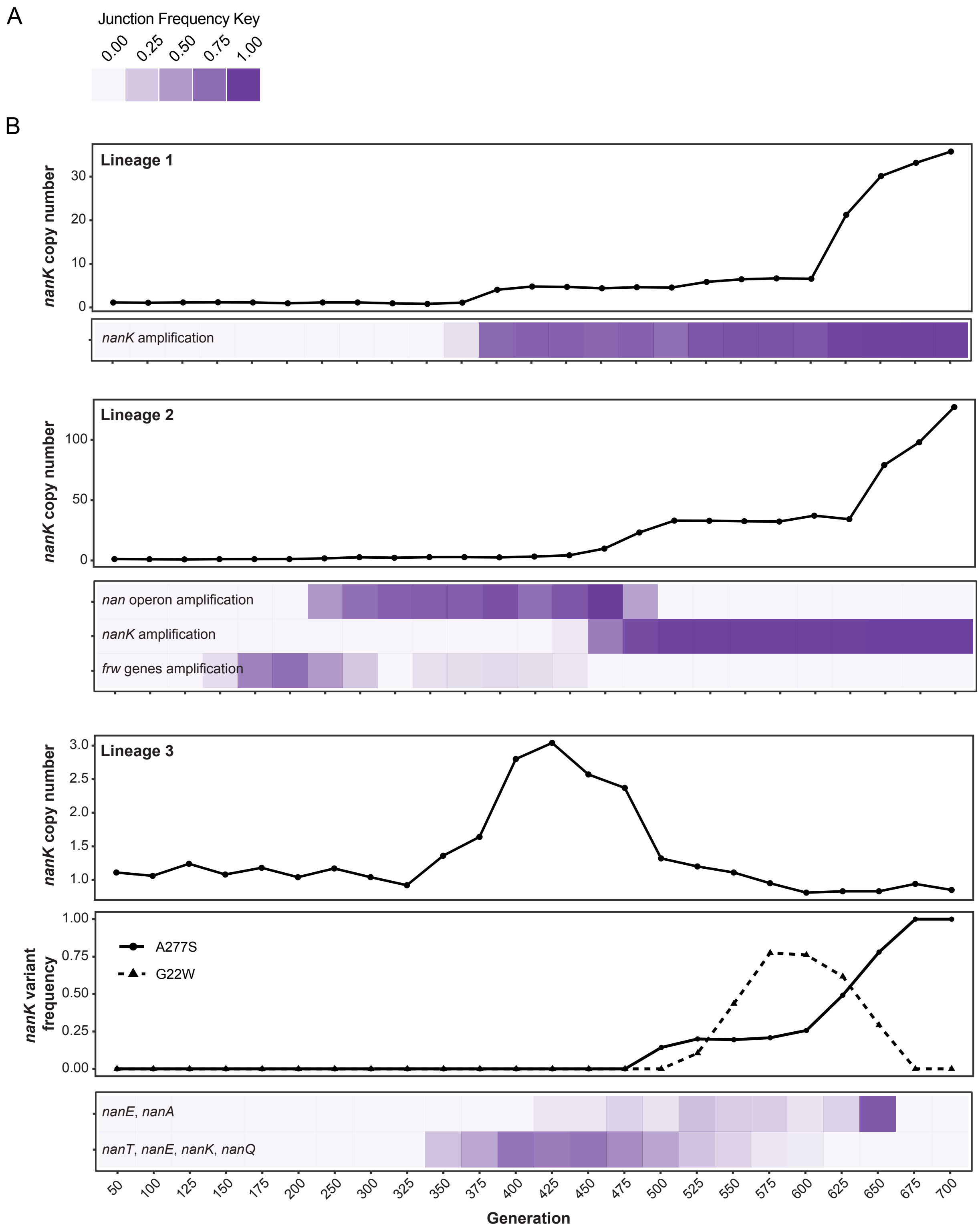


**Figure S4**: Comparison of *nanK* copy number, population frequency of amplifications, and NanK variants. (A) Key relating color opacity and amplification frequency. (B) *nanK*

copy number and amplification frequency for Lineages 1 (top), 2 (middle), and 3 (bottom) throughout the adaptive timeline. The frequency of A277S and G22W NanK variants is shown for Lineage 3.

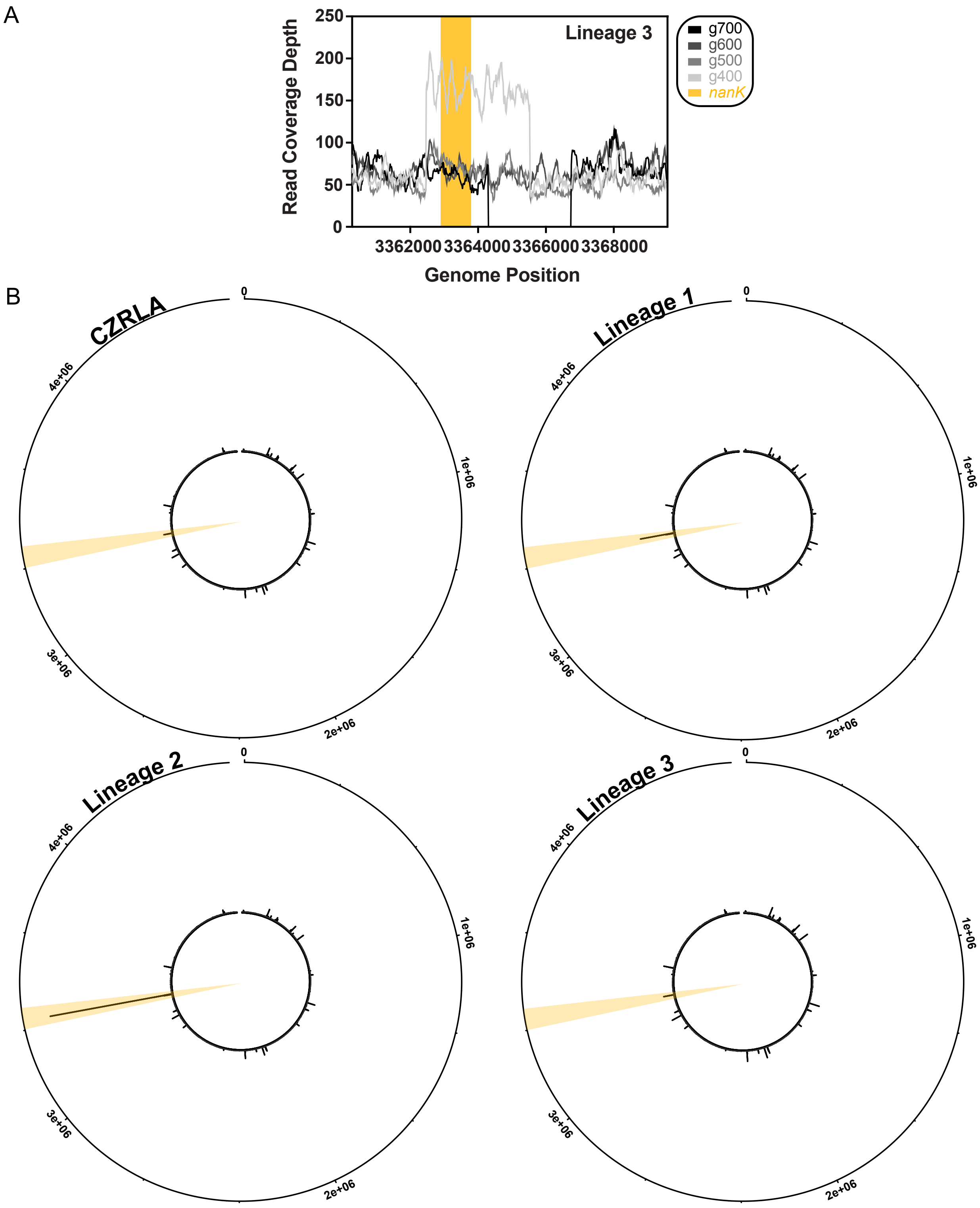


**Figure S5:** Representations of genome coverage from Illumina sequencing. (A) Raw read coverage depth from population sequencing of Lineage 3 cells at generations 400, 500, 600, and 700. Genomic region spans the *nan* operon per genome coordinates of the *E. coli* BW25113 genome (NCBI: CP009273.1). The region encoding *nanK* is yellow. (B) Normalized genome-wide read coverage depth for CZRLA control cells and population samples of Lineages 1, 2, and 3 at generation 700. An approximate region spanning the *nanK* coding sequence is yellow. All plots were made on the same scale, such that a genomic copy number of 10 represents the same line length in all plots.

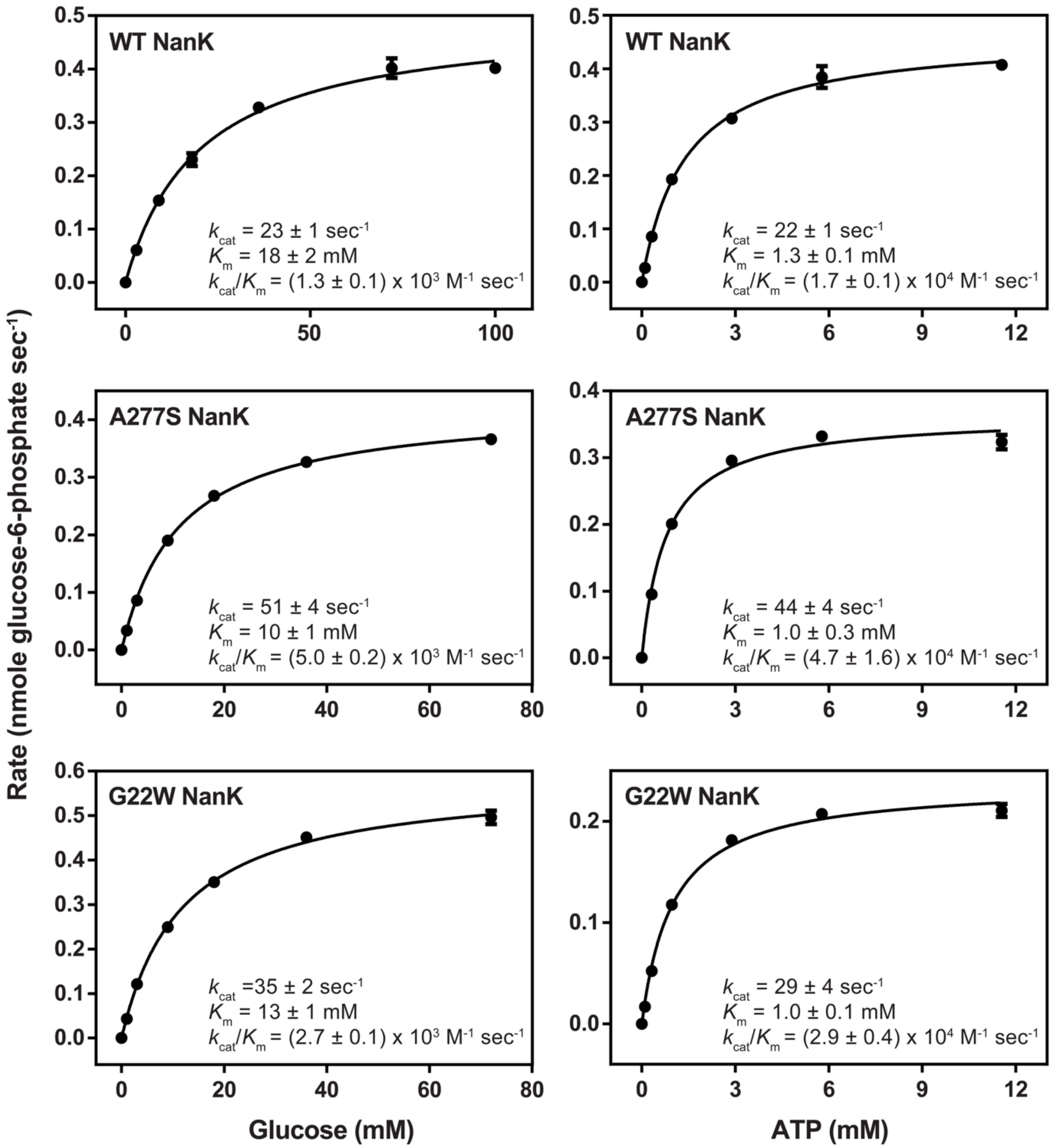


**Figure S6:** Representative replicate kinetic curves used to extract $k_{cat}$ and $K_m$ values for wild-type (top), A277S (middle), and G22W (bottom) NanK catalyzing glucose-6-phosphate synthesis. All data were fitted using the Michaelis-Menten equation. Points represent (N=3) ± SD.

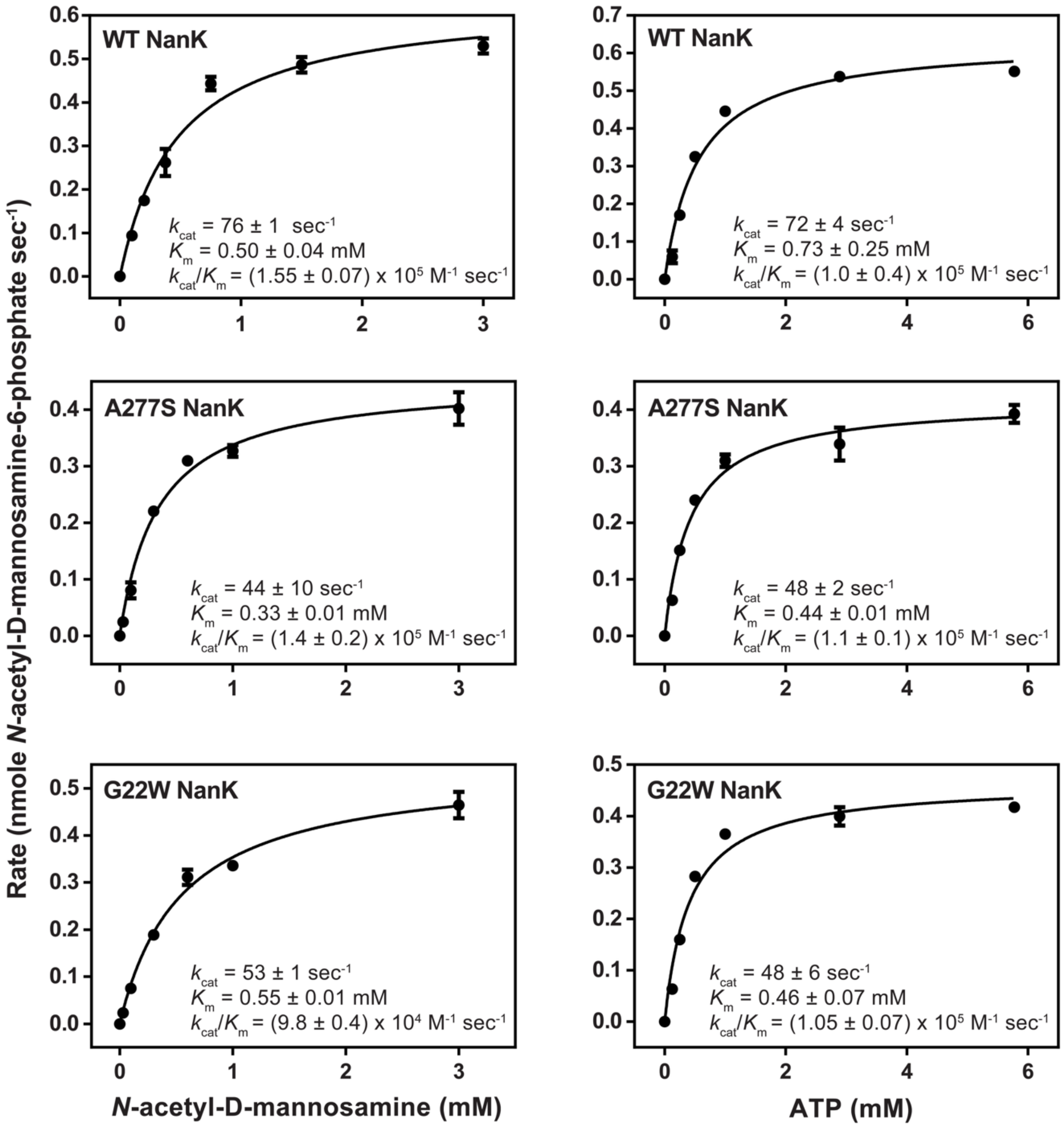


**Figure S7:** Representative replicate kinetic curves used to extract $k_{cat}$ and $K_m$ values for wild-type (top), A277S (middle), and G22W (bottom) NanK catalyzing *N*-acetyl-D-mannosamine-6-phosphate synthesis. All data were fitted using the Michaelis-Menten equation. Points represent (N=3) ± SD.

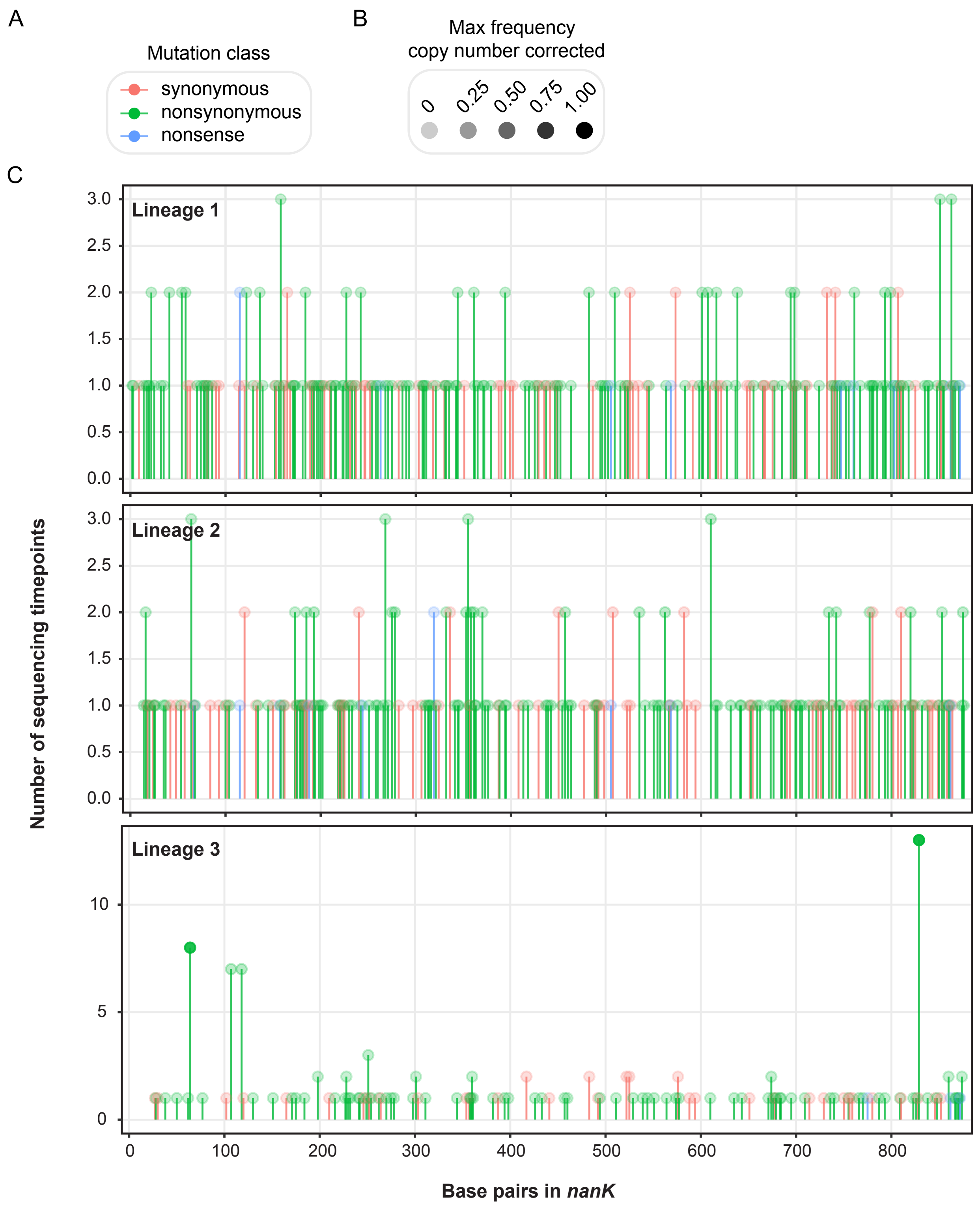


**Figure S8**: Low-frequency alternate base substitutions in the *nanK* coding sequence. (A) Key demonstrating the relationship between mutation class and color. (B) Key relating the opacity of alternate base positions with their maximum frequency in the population. (C) All examples of alternate base substitutions observed in the *nanK* sequence in Lineage 1 (top), 2 (middle), and 3 (bottom). Alternate bases were analyzed from one sequencing timepoint before amplification to the end of ALE for all Lineages.

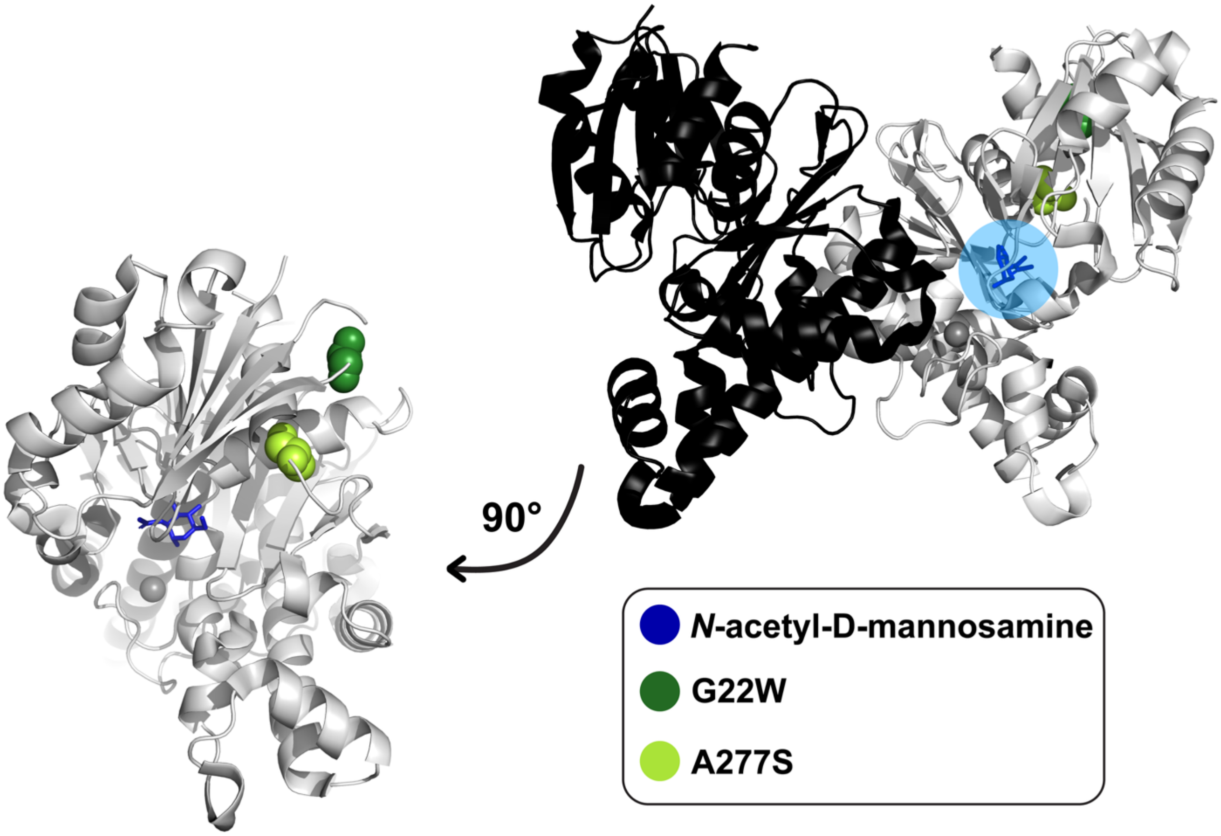


**Figure S9:** Unliganded crystal structure of *E. coli* NanK (PDB 2AA4) with *N*-acetyl-D-mannosamine modeled into one active site (crystal structure of liganded *Haemophilus influenzae*, PDB 6JDC). *N*-acetyl-D-mannosamine is shown as a blue stick representation. Positions G22 and A277 are shown as dark green and light green spheres, respectively.

### Supplementary Results and Discussion: Additional Mutations

#### Miscellaneous mutations common to adaptive laboratory evolution studies appear in all Lineages

During initial medium conditioning, CZRLA cells gained a fixed I774T mutation in *rpoC*, which encodes the β' subunit of RNA polymerase (Figure S2). *rpoC* mutations are common in *E. coli* ALE experiments and are associated with enhanced growth on minimal medium.[2–6] During genetic editing, BW25113 Δ*glk,* Δ*pts* cells developed intergenic mutations between *gltU* and *rrlC, dnaB* and *alr*, and a IS*5* 4 base-pair insertion in the coding sequence of *yobF* (Figure S2). *gltU* encodes a glutamate tRNA and *rrlC* encodes the 23S subunit ribosomal RNA. Mutations in *rrlC* and its transcription unit have been observed in numerous ALE studies.[2,7,8] *dnaB* encodes a replicative DNA helicase and *alr* encodes alanine racemase 1. A similar intergenic mutation was observed during adaptation of *E. coli* to norfloxacin.[7] *yobF* encodes a protein of unknown function, but the same mobile element mutation has been observed in *E. coli* during adaptation to conditions including acid stress, ionic liquids, and alternating carbon sources.[9–11]

In the first 100 generations of adaption, lineages acquire transient mutations in *proQ* (encoding RNA chaperone, ProQ) (Figure S2). ProQ is an RNA-binding protein that interacts with hundreds of transcripts.[12] ProQ truncations have been observed in ALE conditions including adaptation to antimicrobials, alcohols, and commodity chemicals.[7,13–16]

#### Unique mutations in putative *pts* enzymes likely do not directly impact glucose processing

In addition to mutations in expected targets, all Lineages developed mutations that were not observed in previous work.[1] Lineages 1 and 2 developed separate disruptive mutations by generation 50 in *dhaM,* which encodes dihydroxyacetone kinase subunit M (DhaM) (Figure 1). Intriguingly, DhaM bears ~30% sequence identity to two components of the phosphotransferase system (PTS), HPr and EI, which were deleted from our parental strain.[17] While the *dhaM* frameshift in Lineage 1 reaches a maximum frequency of ~18% before exiting the population, the *dhaM* truncation in Lineage 2 appears initially at ~80% and reaches fixation by generation 325. Lineage 2 also demonstrates gene amplification of a region encoding putative PTS enzymes PtsA/FrwA, FrwB, FrwC, and FrwD, as well as putative pyruvate formate lyase PflD and putative pyruvate formate lyase activating enzyme PflC (Figure S10).[18] The *frw* genes encode proteins with homology to a complete fructose PTS, with the exception of an HPr-like enzyme.[18] Amplification of this region occurs from generation 175 to 300 and peaks in frequency at ~74% at generation 200 (Figure S10). The ramifications of mutations in putative PTS enzymes were estimated in two ways. First, mutational timeline and specific activity data were used to approximate which mutations likely contributed to significant changes in total glucokinase or *N*-acetyl-D-mannosamine kinase activity during adaptation (Figures 1, 2A). No Lineage demonstrates significant specific activity changes until generation 500, suggesting that mutations that accumulated before that point, at least in isolation, did not lead to observable changes in kinase activity. All putative PTS or pyruvate formate lyase enzyme mutations appear prior to generation 300, indicating that they do not have an observable effect on phosphorylation of either glucose or *N*-acetyl-D-mannosamine.

To investigate the possibility that the *dhaM* truncation in Lineage 2 caused a reconstruction of the deleted PTS, MacConkey agar was used to screen glucose transport and fermentation (Figure S11). MacConkey agar is a rich differential medium that is used to isolate gram negative bacteria and screen for sugar fermentation.[19] Sugar fermentation will acidify the medium and cause fermenting cells to grow rich pink colonies, due to neutral red indicator in the medium. Otherwise, colonies are a pale yellow.[19] To identify phenotypic changes due to the *dhaM* mutation, Lineage 2 cells were compared between generations 0 and 50, as the only significant mutation developed between these generations is the *dhaM* truncation mutation at a population frequency of ~0.8 (Figure 1). Colonies of Lineage 2 cells were pale yellow on glucose MacConkey agar at generations 0 and 50, indicating that the high frequency *dhaM* mutation does not cause a detectable change in fermentation activity. However, generation 50 cells have pink colonies when grown on *N*-acetyl-neuraminic acid (Neu5Ac) (Figure S11). Thus, these mutations may contribute to utilization of pyruvate, which is generated through cleavage of Neu5Ac into *N*-acetyl-D-mannosamine. Deactivation of *dhaM*, either through a frameshift or truncation mutation, would prevent DhaM from transforming pyruvate, which is the product of its native reaction. This is supported by pink colony formation on MacConkey agar when cells with the *dhaM* truncation are grown on Neu5Ac (Figure S11). The putative pyruvate formate lyase enzymes may process pyruvate, which is generated from cleavage of Neu5Ac to *N*-acetyl-D-mannosamine, to acetate.

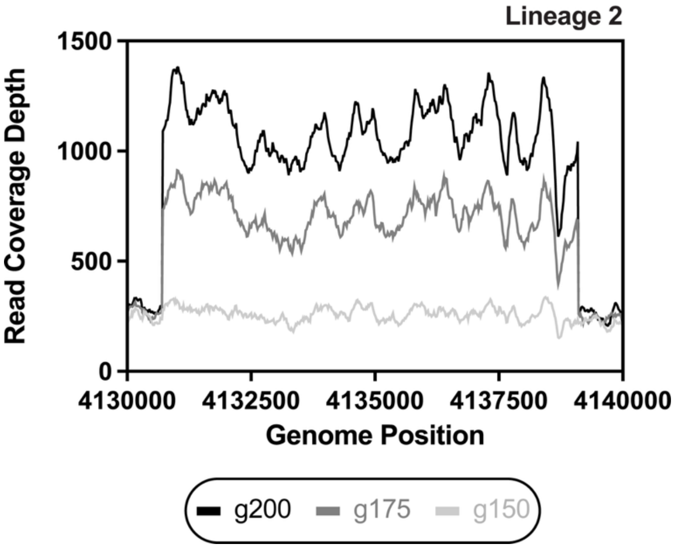


**Figure S10:** Raw read coverage depth data showing amplification of putative phosphotransferase system and pyruvate formate lyase enzymes in Lineage 2. The displayed genome positions include the loci of *frwA, frwB*, *frwC*, *frwD*, *pflC*, and *pflD* based on the *E. coli* BW25113 genome (NCBI CP009273.1). Read coverage depth is represented in solid, grayscale lines for generations 150, 175, and 200.

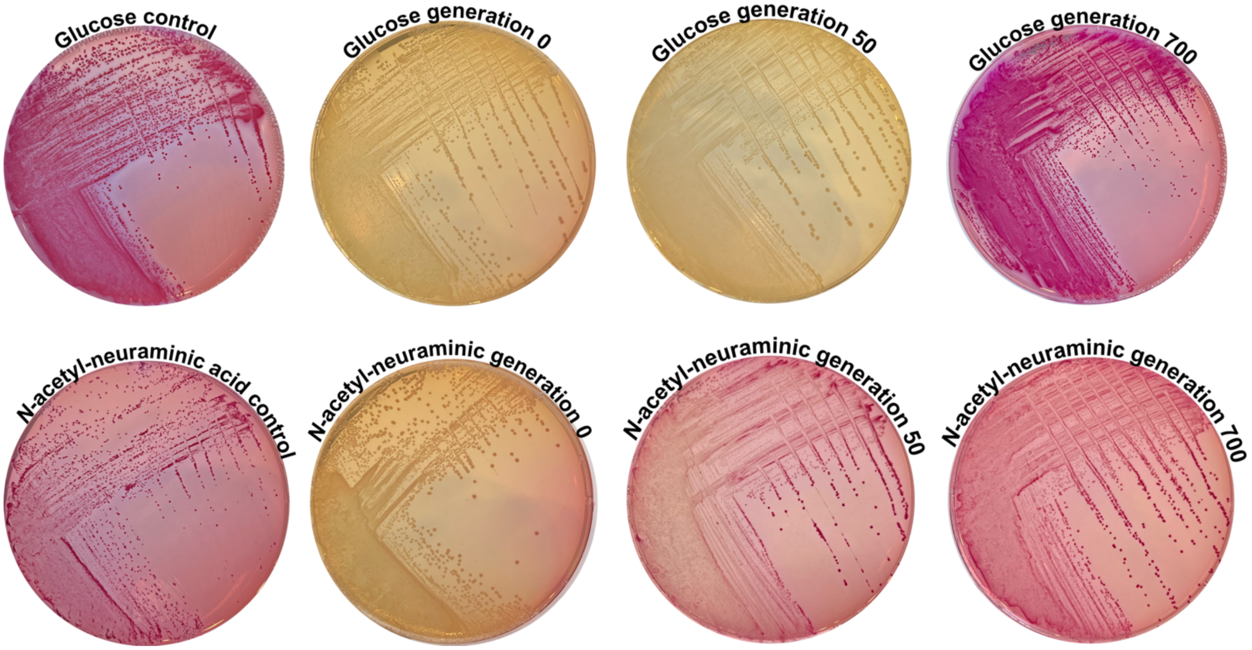


**Figure S11:** MacConkey agar test to investigate the effect of *dhaM* truncation in Lineage 2 cells throughout adaptation. Strain CZRLA was used as a control for plating on MacConkey agar containing an equivalent concentration of either glucose (top) or Neu5Ac (bottom). Lineage 2 cells are shown following growth on MacConkey agar supplemented with either substrate at generations 0, 50, and 700. Generation 50 cells contain a *dhaM* truncation at a population frequency of ~0.8 (Figure 1).

### Materials and Methods

#### *Fitness assay of cells grown on Neu5Ac*

A strain of BW25113 *E. coli* that was previously adapted to grow on 0.2% glucose M9 minimal medium was streaked onto an LB agar plate and grown overnight.[1] Three colonies were selected as biological replicates. Each colony was inoculated into 10 mL of LB medium and grown with shaking overnight. Six 1 mL aliquots of overnight culture were removed, pelleted, and washed three times with M9 minimal medium containing 0.2% glucose and either 0.5 mM, 1 mM, 3 mM, 7 mM, 11 mM, or 20 mM Neu5Ac. Washed cultures were plated in a sterile 96-well microplate (Corning) at a 1:250 dilution into fresh M9 medium of the appropriate condition. The plate was covered with breathable film (Diversified Biotech) and incubated in a Spectramax iD5 Multi-Mode microplate reader at 37°C with shaking. $OD_{600\ nm}$ measurements were taken every 15 minutes to monitor growth. Following 6 hours of incubation, cultures were again diluted 1:250 into fresh M9 medium in a new 96-well microplate and incubated as described. Cultures were incubated until all samples reached saturation. Growth rates were extracted from $OD_{600\ nm}$ measurements using the Amiga program.[20]

#### *Glucose and Neu5Ac medium conditioning*

Adaptation of *E. coli* BW25113 to the M9 minimal medium conditions used during ALE was performed using standard methods.[21] Three independent biological replicates were seeded from each of three colonies of *E. coli* BW25113 cells and serially passaged on M9 minimal medium containing 11 mM glucose and 3 mM Neu5Ac for 500 generations. Cells were passaged once per day during stationary growth phase. Following 500 generations of growth, the fitness of medium-adapted cells was compared to wild-type BW25113 in a fitness assay as previously described. The biological replicate that demonstrated the shortest lag time was selected as the parental strain for genetic editing and ALE. This strain was named CZRLA.

#### *Genetic editing*

CZRLA cells were made glycoslysis-deficient through deletions of the *glk* and *ptsH*, *ptsI*, and *crr* genes using λ red recombineering.[22] CZRLA cells were made electrocompetent and transformed with the helper plasmid pSIJ8, which harbors λ red proteins gamma, beta, and exonuclease under the control of an arabinose-inducible promoter, and flippase recombinase under the control of a rhamnose-inducible promoter.[22] Cells with pSIJ8 were grown to an $OD_{600\ nm}$ of 0.3, then λ protein expression was induced with 15 mM arabinose. Cultures were grown until they reached an $OD_{600\ nm}$ of 0.6, then growth was quenched on ice and cells were made electrocompetent. A kanamycin resistance gene was PCR amplified from plasmid pKD4 using primers that added on homology arms for the *pts* operon locus.[23] 200 ng of the *pts*-targeting kanamycin resistance cassette was transformed into electrocompetent cells expressing λ red proteins. Cells were recovered, then streaked onto LB agar plates (35 µg/mL kanamycin) to select for transformants with the kanamycin gene integrated into their genomes. Integration of the kanamycin resistance gene into the *pts* operon locus was verified through colony PCR on the *pts* locus. Δ*pts::kanR* + pSIJ8 cells were grown to an $OD_{600\ nm}$ of ~0.1, then expression of flippase recombinase was induced with the addition of 50 mM rhamnose. Cells were incubated for an additional 4 hours. The removal of the kanamycin resistance gene from the *pts* locus was verified by lack of growth on medium containing kanamycin and by colony PCR of the *pts* locus. This process was repeated in the Δ*pts* strain to delete the *glk* gene, yielding the parental ALE strain BW25113 Δ*pts* Δ*glk*. Plasmid pSIJ8 was cured from the strain following growth at 37°C. Deletion of the *pts* operon and *glk* gene were verified using Sanger sequencing. Primer sequences used for λ red recombineering and colony PCR are listed in Table S3.

***Adaptive laboratory evolution***

BW25113 Δ*pts* Δ*glk* cells were grown overnight on LB-agar plates. To initiate replicate lineages, three colonies were inoculated into 50 mL aliquots of LB medium and grown with shaking at 37°C to an approximate $OD_{600\ nm}$ of 1.0. Each culture was removed from incubation and growth was quenched on ice. Cultures were pelleted and washed 3 times with M9 minimal medium containing 0.2% glucose and 3 mM Neu5Ac. Cultures were inoculated into each of 3 aliquots of 500 mL M9 minimal media containing 0.2% glucose and 3 mM Neu5Ac to an approximate initial $OD_{600\ nm}$ of 0.06 and grown with shaking at 37°C. The remaining culture from each replicate was used to make permanent frozen stocks. Lineages were cultured through serial passage by transferring the evolving replicate populations at an $OD_{600\ nm}$ of approximately 1.0. Lineages were transferred into fresh aliquots of selective medium at an initial $OD_{600\ nm}$ of approximately 0.03 (~$1.2 \times 10^{10}$ cells to seed each new culture flask). Cells were cultured in this manner in M9 minimal medium containing 0.2% glucose and diminishing concentrations of Neu5Ac acid. Neu5Ac was provided at 3 mM up to generation 100, 1.5 mM until generation 300, 0.75 mM until generation 500, 0.375 mM until generation 600, and 0 mM until generation 700. At the time of each transfer, permanent frozen stocks were made of each replicate culture.

***Illumina whole genome sequencing and Breseq polymorphism detection***

Genomic DNA was extracted from frozen stocks of cultures made throughout ALE using the EZNA Bacterial DNA Kit (Omega Bio-Tek). DNA libraries were constructed using the NEBNext Ultra II DNA Kit and sequenced on a NOVASEQ 6000 or NOVASEQ X Plus sequencing system. Resulting whole genome sequences were aligned to the reference BW25113 genome using Breseq to identify polymorphisms.[24] Per standard Breseq usage, a minimum frequency threshold of 5% was used in our analysis. For simplicity, and to minimize discussion of mutations that are unlikely to impact cellular fitness, only mutations that reached at least 10% frequency in the population and were present for at least two consecutive sequencing timepoints are discussed in this manuscript.

***Fitness assays on adapted Lineages***

CZRLA, BW25113 Δ*glk* Δ*pts* (L84P NanK), BW25113 Δ*glk* Δ*pts* (L84R NanK), and BW25113 Δ*glk* Δ*pts* ALE Lineage strains were inoculated from frozen stocks into 5 mL of LB.[1] Populations were sampled directly from frozen stocks to sample the diversity of cell genotypes at each time point. Cultures were grown at 37°C until they reached an $OD_{600\ nm}$ of 1-2. Incubation was halted, total cell count was normalized, and cells were washed three times with the appropriate selective medium (M9 minimal medium containing either 11 mM glucose or 3 mM Neu5Ac). Each population was diluted into fresh medium to an initial $OD_{600\ nm}$ of 0.2. Populations were plated in triplicate into a 96-well plate (Corning) and sealed with a breathable membrane (Diversified Biotech). Cells were incubated at 37°C with shaking in a BMG Labtech multi-mode microplate reader. An absorbance measurement at 600 nm was recorded every 15 minutes until samples reach saturation. Growth rates and lag times were extracted from the $OD_{600\ nm}$ measurements using the AMiGA program.[20]

***Total glucokinase and N-acetyl-D-mannosamine kinase activity in whole cell lysates***

CZRLA and ALE replicate Lineage cells from generations 100, 300, 500, 600, and 700 were grown overnight in LB. Following outgrowth, cells were washed three times in M9 minimal medium. The carbon source(s) in the medium reflected that of the medium conditions in which adapting cells were initially cultured, i.e., cells at generation 700 were grown in solely 11 mM glucose. CZRLA cells were grown in either 3 mM Neu5Ac and 11 mM glucose or solely 11 mM glucose. Washed cells were inoculated 1:100 into 100 mL of fresh M9 minimal medium and grown at 37°C with shaking to an approximate $OD_{600\ nm}$ of 1 in biological triplicate for each strain. Cells were harvested by centrifugation at 6,000 x g for 10 minutes at 4°C, supernatant was removed, and cell pellets were frozen at -20°C. Cell pellets were resuspended in 1 mL of cold buffer containing Tris (pH 7.6, 25 mM), $MgSO_4$ (5 mM), and glycerol (5%). Cells were lysed by sonication and cellular debris was pelleted by centrifugation at 15,557 x g for 30 minutes at 4°C. Supernatants containing clarified cell lysates were saved. Total protein concentration was determined using a BCA assay (Pierce).

Total glucokinase activity in cell lysates was determined by coupling the oxidation of NADP+ to the production of glucose-6-phosphate through glucose-6-phosphate dehydrogenase as previously described.[25] Assays contained Tris (pH 7.6, 200 mM), DTT (1 mM), $MgCl_2$ (27 mM), NADP+ (0.5 mM), glucose-6-phosphate dehydrogenase (15 U), and saturating concentrations of glucose (290 mM) and ATP (27 mM). Assays were performed at 30°C and initiated through addition of glucose.

Total *N*-acetyl-D-mannosamine kinase activity in cell lysates was determined by coupling the oxidation of NADH to the production of ADP through the combined activity of pyruvate kinase and lactate dehydrogenase as previously described.[25] Assays contained Tris (pH 7.6, 200 mM), DTT (1 mM), $MgCl_2$ (13.5 mM), NADH (0.5 mM), KCl (5 mM), phosphoenol pyruvate (5 mM), pyruvate kinase (40 U) and lactate dehydrogenase

(20 U) and saturated concentrations of *N*-acetyl-D-mannosamine (5 mM) and ATP (13.5 mM). Assays were conducted at 30°C and initiated by addition of *N*-acetyl-D-mannosamine.

All assays were conducted in biological triplicate for each ALE Lineage. Each biological replicate was measured in technical duplicate or triplicate and specific activity was averaged between 6 to 9 individual data points per ALE Lineage. Specific activity is reported as the average activity for all technical and biological replicates per ALE Lineage in μmol enzyme $min^{-1}$ mg $protein^{-1}$. Specific activity of CZRLA lysates following cell outgrowth on 11 mM glucose and 0 mM Neu5Ac was also determined, but *N*-acetyl-D-mannosamine kinase activity was undetectable under these conditions, and therefore is not used for comparison.

***RT-qPCR to detect transcriptional changes in all candidate kinases***

Cells from all ALE Lineages at generations 100, 350, and 700 were inoculated into 5 mL of LB medium from frozen stocks. Cells were grown overnight at 37°C with shaking. Following outgrowth, 1 mL of each culture was pelleted and washed three times with selective M9 medium. For each Lineage at each generation, three independent cultures were inoculated 1:1000 into M9 medium. Medium conditions represented the culture conditions of when cells were initially grown, that is, 11 mM glucose and 3 mM Neu5Ac for generation 100 cells, 11 mM glucose and 0.75 mM Neu5Ac for generation 350 cells, and 11 mM glucose for generation 700 cells. Each of these cultures was grown at 37°C with shaking to an approximate $OD_{600\ nm}$ of 1.0, then cells were harvested and made into permanent frozen stocks. For controls, CZRLA cells were grown in the same manner in 11 mM glucose M9 minimal medium either with or without 3 mM Neu5Ac.

RNA was extracted from permanent frozen stocks using the Omega BioTek E.Z.N.A. Bacterial RNA Kit (Omega BioTek, Norcross, GA). 5 μg of RNA was DNAse treated twice with DNAse I RNAse free from NEB following manufacturer's instructions (NEB, Ipswich MA). DNAse treated RNA was used for cDNA synthesis using SuperScript III First-Strand synthesis kit (Thermo Fisher Scientific, Waltham, MA). cDNAs were used as a template for qPCR on a QuantStudio 7 Flex real-time PCR System (ThermoFisher Scientific) using PerfeCTa SYBR® Green SuperMix, Low Rox (Quanta Bioscience). Fold expression was determined by normalization to the endogenous housekeeping gene *cysG*.[26] Primers used in this study were chemically synthesized by Eurofins Genomics LLC (Louisville, KY) and are listed in Table S4. Relative gene expression was calculated from cycle threshold values using the $2^{-\Delta\Delta CT}$ method.[27]

#### *MacConkey agar assay to test dhaM effect*

CZRLA, ALE Lineage 2 (generation 0), ALE Lineage 2 (generation 50), and ALE Lineage 2 (generation 700) cells were spread-streaked onto MacConkey agar (Difco) plates from permanent frozen stocks. For each strain, cells were plated onto MacConkey agar containing 32 mM glucose and MacConkey agar containing 32 mM Neu5Ac. Plates were incubated at 30°C until robust colony formation was observed (Figure S11).

#### *Gene amplification analysis on Illumina whole genome sequencing*

Average gene copy numbers throughout ALE were estimated from Illumina whole genome sequencing population data (whole genome sequencing described above). BAM files generated by Breseq during mapping of sequencing reads to the BW25113 reference genome (NCBI CP009273.1) were used to investigate copy number. Per sample, the average read coverage depth across each gene in the BW25113 genome (NCBI CP009273.1) was divided by the average read coverage depth of the entire genome to investigate the relative coverage level). This process was also performed for CZRLA control cells. To visualize changes in read coverage depth, raw read coverage depth per genome coordinate was plotted over time for amplified regions of interest. To investigate if changes in read coverage depth represented tandem duplications and/or translocations, all reads partially aligning to the *nanK* locus were compared through an NCBI nucleotide blast to the BW25113 genome to investigate read alignment. Gene copy number was plotted in circular genome plots to visualize genomic changes in copy number relative to the parental strain (Figure S5).

"New junction evidence" reported from Breseq data was used to determine the population frequency of amplifications for each Lineage across ALE (Figure S4).[24,28,29] The genome coordinates of gene amplifications were compared with those in new junction evidence to confirm their connection.

Following initial Breseq analysis, BAM files were used to further investigate possible instances of mutations in *nanK* copies at low frequency. This is because Breseq analysis intrinsically filtered out polymorphisms that appeared below 5% in the populations, and Breseq is agnostic to changes in copy number. For example, consider if 100% of a population had 30 copies of *nanK*, and 1/30 of those copies had a fixed mutation in the *nanK* coding sequence. Despite fixation of that mutation in one of the copies, without accounting for multiple copies, mutation frequency would be counted as only 3.3% of cumulative *nanK* reads. Therefore, we suspected mutations in *nanK* may not have been reported through initial Breseq analysis on Lineages 1 and 2 due to artificial dilution of mutation frequency resulting from amplification.

To this end, BAM files from Breseq were used to identify all bases that aligned to any position in the *nanK* reference sequence. If a sequenced base differed from the reference, we calculated the frequency of an alternate base at that position by dividing the number of reads where the read base pair deviated from the reference by the total number of reads that aligned at that specific position. Using the *nanK* copy number estimates calculated previously, alternate base frequencies were corrected for *nanK* copy number dilution by multiplying the frequency by the estimated *nanK* copy number at that sequencing timepoint. We performed this analysis for all Lineages starting from one sequencing time point before the onset of amplification until the end of ALE.

Any position for which an alternate base was called was analyzed for its persistence throughout adaptation. Any position for which an alternate base was called was also classified as a nonsynonymous, synonymous, or nonsense mutation (Figure S8).

***ONT sequencing on Lineage 1 and 2 cells***

Lineage 1 (generation 700) and Lineage 2 (generation 700) cells were spread-streaked onto LB agar plates from permanent frozen stocks. Plates were incubated at 37°C overnight. Two individual colonies per plate were inoculated into LB medium and incubated at 37°C with shaking overnight. Genomic DNA was extracted from overnight cultures using the EZNA Bacterial DNA Kit (Omega Bio-Tek). ONT long-read sequencing was performed by Plasmidsaurus. This process was repeated with cells grown on 0.2%

glucose M9 minimal medium rather than LB. Three clones per Lineage were grown on M9 minimal medium, yielding 5 total clones studied per Lineage.

Reads containing *nanK*-like sequences were identified through an NCBI nucleotide blast of all raw ONT reads against the *E. coli* BW25113 *nanK* coding sequence (NCBI: CP009273.1). Blast results were filtered by length to exclude partial alignment to *nanK* (size cut-off = 0.8**nanK* length). Raw data and reads identified through blast against *nanK* were inspected for general quality using NanoPlot.[30] Reads containing *nanK*-like sequences were aligned to the BW25113 genome using Minimap2 and visualized with IGV.[31,32] Reads containing *nanK*-like sequences were also blasted against the BW25113 genome to investigate the identity of sequences flanking *nanK*. Nucleotide blasts of reads revealed that *nanK* repeats were consecutive and non-overlapping, indicating each *nanK* read represented an individual repeat.

#### *Expression and purification of NanK mutants*

Wild type, A277S, and G22W NanK variants were expressed from pCA24N vectors. A277S and G22W NanK mutants were created from the wild type pCA24N-*nanK* vector using Q5 site-directed mutagenesis.[33] Both NanK mutants were expressed in *E. coli* BW25113 Δ*glk* Δ*pts* cells to prevent contamination from native glucokinases.[16] *E. coli* BW25113 Δ*glk* Δ*pts* cells with pCA24N-(A277S *nanK*) or pCA24N-(G22W *nanK*) expression plasmids were grown overnight at 37°C in LB medium containing 25 μg/mL chloramphenicol. 2 mL of each overnight culture were inoculated into 1L of LB medium containing 25 μg/mL chloramphenicol. Cells were grown at 37°C with shaking to an approximate $OD_{600\ nm}$ of 0.5, then protein expression was induced with IPTG (final concentration 1 mM). Incubator temperature was reduced to 20°C and cells continued to grow overnight. Cell pellets were harvested through centrifugation at 6000 x g and 4°C for 10 minutes, then frozen at -20°C.

Cell pellets were resuspended in buffer containing sodium phosphate (20 mM, pH 7.4), imidazole (30 mM), and NaCl (500 mM). Cells were lysed by sonication. Cellular debris was separated through centrifugation at 15,600 x g at 4°C for 30 minutes. Clarified lysates were loaded onto a HisTrap FF crude column that had been washed with resuspension buffer. NanK mutants were eluted from columns in buffer containing sodium phosphate (20 mM, pH 7.4), imidazole (300 mM), and NaCl (500 mM). Purified NanK mutants were dialyzed against two sequential 1L aliquots of Tris (25 mM, pH 7.6) containing $MgSO_4$ (5 mM), glycerol (5%), and DTT (0.5 mM). Protein concentrations were estimated spectrophotometrically using absorbance at 280 nm and a molar extinction coefficient of 15,000 $M^{-1}$ $cm^{-1}$ as described previously.[25] The purity of all enzymes was estimated to be >95% through polyacrylamide gel electrophoresis.

***Steady state kinetic assays of NanK mutants***

A277S and G22W NanK were purified as described above. Glucokinase activity was measured by coupling the production of glucose-6-phosphate to the reduction of NADP+ with glucose-6-phosphate dehydrogenase, as previously described.[25] Assays contained Tris (0.2 M, pH 7.6), NADP+ (0.5 mM), DTT (1 mM), ATP (12 mM), $MgCl_2$ (12 mM), and glucose-6-phosphate dehydrogenase (7.5 units). The glucose concentration was varied between 1 and 72 mM. The apparent ATP $K_m$ was determined under the same conditions, but with glucose held at saturating concentration (72 mM) and ATP was varied between 0.1 or 0.32 and 11.6 mM. $MgCl_2$ concentration was held 1:1 with ATP. Assays were performed at 25°C and initiated with the addition of enzyme. Reaction rates were determined by measuring the change in absorbance at 340 nm.

*N*-acetyl-D-mannosamine kinase activity was measured by coupling the production of ADP with the oxidation of NADH through pyruvate kinase and lactate dehydrogenase, as previously described.[25] Assays contained Tris (0.2 M, pH 7.6), NADH (0.5 mM), KCl (5 mM), DTT (1 mM), phosphoenol pyruvate (5 mM), ATP (5.6 mM), MgCl2 (5.6 mM), pyruvate kinase (20 U), and lactate dehydrogenase (10 U). *N*-acetyl-D-mannosamine concentration was varied between 3 and 0.03 mM. The apparent ATP $K_m$ was determined under the same conditions, but with *N*-acetyl-D-mannosamine held at saturating concentration (3 mM) and ATP was varied between 5.6 and 0.125 mM. $MgCl_2$ concentration was held 1:1 with ATP. Assays were performed at 25°C and initiated with the addition of *N*-acetyl-D-mannosamine. Reaction rates were determined by measuring the change in absorbance at 340 nm.

**Table S3: Primers used during genetic editing**

| | Primer | Primer Sequences 5'→3' |
|---|---|---|
| Colony PCR | *glk* F | TCGACCTTGTGAGGGGACAC |
| | *glk* R | CTGCAATTGGTGCTGAAACG |
| | *pts* F | TATTGCGCTCTTCGTGCGTC |
| | *pts* R | GCTACACCCAGCAGCATGAG |
| λ red | *glk* F | CCCAGGTATTTACAGTGTGAGAAAGAATTATTTTGACTTTAGC GGAGCAGTTGAAGAATGGTGTAGGCTGGAGCTGCTTC |
| | *glk* R | ATTATCGGGAGAGTTACCTCCCGATATAAAAGGAAGGATTTAC AGAATGTGACCTAAGGTGGTCCATATGAATATCCTCC |
| | *pts* F | CTAGACTTTAGTTCCACAACACTAAACCTATAAGTTGGGGAA ATACAATGGTGTAGGCTGGAGCTGCTTC |
| | *pts* R | ATGGGCGCCATTTTTCACTGCGGCAAGAATTACTTCTTGATG CGGATAACGGTCCATATGAATATCCTCC |

**Table S4: Primer sequences used for RT-qPCR**

| GENE | SEQUENCE 5' → 3' |
|---|---|
| *alsK* F | GGCAGGTTAGGCGTAGAAATA |
| *alsK* R | GCAACTCAGGCGCTTTAAC |
| *cysG* F | AGGTGGCGATCCGTTTATTT |
| *cysG* R | AATAGGCAGAGCAACCAGAAG |
| *mak* F | GCGAGGAAGTCCCTTGTTATT |
| *mak* R | GTCCGCTCAAACGACGATAA |
| *nagK* F | CGCGATGTACGCCTTGATAA |
| *nagK* R | CCCATCACCAGTGGATATTGAG |
| *nanK* F | CCGGTGGAAACGGTGATAAA |
| *nanK* R | TTGCCGACCATTGCCATTA |
